# Filter-aided extracellular vesicle enrichment (FAEVEr) for proteomics

**DOI:** 10.1101/2023.07.06.547926

**Authors:** Jarne Pauwels, Tessa Van de Steene, Jana Van de Velde, Freya De Muyer, Danaë De Pauw, Femke Baeke, Sven Eyckerman, Kris Gevaert

## Abstract

Extracellular vesicles (EVs), membrane-delimited nanovesicles that are secreted by cells into the extracellular environment, are gaining substantial interest due to their involvement in cellular homeostasis and their contribution to disease pathology. The latter in particular has led to an exponential increase in interest in EVs as they are considered to be circulating packages containing potential biomarkers and are also a possible biological means to deliver drugs in a cell-specific manner. However, several challenges hamper straightforward proteome analysis of EVs as they are generally low abundant and reside in complex biological matrices. These matrices typically contain abundant protein concentrations that vastly exceed those of the EV proteome. Therefore, extensive EV isolation and purification protocols are imperative and many have been developed, including (density) ultracentrifugation, size-exclusion and precipitation methods. Here, we describe filter-aided extracellular vesicle enrichment (FAEVEr) as an approach based on 300 kDa MWCO filtration that allows the processing of multiple samples in parallel within a reasonable timeframe and at moderate cost. We demonstrate that FAEVEr is capable of quantitatively retaining EV particles on filters, whilst allowing extensive washing with the mild detergent TWEEN-20 to remove interfering non-EV proteins. The retained particles are directly lysed on the filter for a complete recovery of the EV protein cargo towards proteome analysis. Here, we validate and optimize FAEVEr on recombinant EV material and apply it on conditioned medium as well as on complex serum. Our results indicate that EVs isolated from MCF7 cells cultured with or without serum have a drastic different proteome because of nutrient deprivation.

## A. Background

Extracellular vesicles (EVs) make up a heterogeneous population of membrane-enclosed particles that are released by virtually all cells into their extracellular (EC) environment and in bodily fluids, such as plasma, urine and cerebrospinal fluid. In general, EVs are subdivided in three main categories according to their biogenesis and size (1). The smallest (30 – 150 nm), termed exosomes, are generated by invaginations at the endosomal membrane, in a transport (ESCRT) dependent (2,3) or independent manner (4). During this process, early endosomes mature into a multivesicular body (MVB) that holds multiple membrane delineated vesicles termed intraluminal vesicles (ILVs). Once the MVB fuses with the plasma membrane (PM), these ILVs are released into the EC environment as exosomes. Microvesicles (100 – 500 nm) are formed by an immediate outward budding of the PM, orchestrated by the ESCRT machinery (5). These protrusions are ultimately pinched off and released from the cell. Finally, apoptotic bodies (50 – 5,000 nm) are formed during apoptotic cell disassembly with large extrusions at the PM, fragmenting dying cells in large blobs that may even contain organelles (6). Ultimately, in the body, different EV categories are released from millions of individual cells, forming an immensely heterogeneous EV pool. This also results in the loss of information on their biogenesis and therefore, in practice, EVs are distinguished from one another by their size; small (sEV, < 200 nm) or large (lEV, > 200 nm), as described in the MISEV2018 guidelines (7). Note that since there is a certain level of size overlap between the different subtypes, none of these subpopulations is exclusive for either the small or the large EV group (8).

Due to their biogenesis, EVs have some important biophysical properties. First, EVs are delineated by a phospholipid bilayer, effectively shielding their cargo (consisting of proteins, oligonucleotides (RNA & DNA), lipids and metabolites) from the hostile extracellular environment, thereby, maintaining the functionality and stability of biomolecules (9). Therefore, the cargo can be considered to represent the molecular state of the parental cell at the moment of EV formation. In addition, (trans)membrane proteins are incorporated into the lipid bilayer, which, in turn, allows targeted signal transduction with distant EV receiving cells and/or allows the selective delivery of their cargo in such cells (10,11). In physiological conditions, this cargo takes part in cellular communication and homeostasis (12,13). In disease, however, the EV cargo can contribute to the pathology, for example by initiating a pre-metastatic niche at a distant location during cancer progression (14,15).

Both of these EV characteristics are appealing for clinical implementations, offering great potential for both therapeutic applications (i.e. targeted drug delivery) as well as for biomarker discovery (e.g. disease diagnosis, prognosis and therapy guidance) (16–18). Although the therapeutic applications remain rather hypothetical or, at best, experimental to date (19,20), the biomarker potential of circulating EVs has known a substantial increase in the past decade. Indeed, EVs are present in different biofluids, such as plasma and urine, which can be isolated in a low to non-invasive way at multiple time points.

In general, considering the biophysical properties of EVs, the identification and characterization of protein biomarkers within or on circulating EVs from liquid biopsies might lead to (early) disease diagnosis, monitoring of disease progression and therapeutic response, all critical aspects towards the establishment of precision treatment (21). In this respect, EVs have shown clinical potential as a source of circulating biomarkers for different cancers (e.g. breast (18), urinary tract (22) and bone cancer (23)) and other diseases such as Alzheimer’s disease (24,25), chronic liver disease (26) and multiple sclerosis (27).

Unfortunately, multiple challenges complicate EV protein research, especially with regard to proteome analysis by bottom-up mass spectrometry (MS), which is the method-of-choice for unbiased protein biomarker discovery (28). Indeed, once EVs are released by the cell they form a heterogeneous pool that resides in biological complex matrices that contain proteins at levels that exceed those of the low abundant EV proteins by several orders of magnitude (29), such as albumin in plasma and uromodulin in urine. In addition, they co-exist with non-EV particles that overlap in size and density such as apolipoprotein particles. As a result, an adequate EV enrichment and purification strategy remains essential for in-depth EV-associated biomarker discovery (30). Of note, in MS-based proteomics, the recent advent of data-independent acquisition (DIA) MS due to computational (31,32) and instrumental (33) improvements has proven to cope more successfully with the above-mentioned challenges compared to traditional data-dependent acquisition (DDA), as it is less biased to relative high protein levels.

Over the years, different EV enrichment strategies have been developed to enrich, purify and concentrate EVs for in-depth proteome analysis and quantitative profiling, with the most prominent strategies being based on size (ultracentrifugation and size-exclusion chromatography) (34,35), density (density gradient ultracentrifugation) (36), precipitation (37,38) or capture of molecular markers present on the EV surface (immuno-capture) (39). To further enhance the purity of EV preparations, different orthogonal approaches can be combined (40,41). However, a golden standard is yet to be determined, as the different strategies not only differ in the consumption of resources and time, but also in terms of effectiveness and reproducibility. Indeed, the different EV isolation techniques introduce variation between experiments in terms of EV count and heterogeneity of molecular (protein) cargo. This has been emphasized in different studies in which EV enrichment strategies were compared (42–46). Along this line, initiatives such as MISEV and EV-TRACK were established within the EV research community to encourage transparency on experimental procedures used to enrich EVs and on acquired results, thereby enhancing inter-lab reproducibility (7,47). Importantly, purified, recombinant EVs are now used as biological reference material to monitor technique development and reproducibility, and improve data normalization (48,49). However, all enrichment strategies have one major bottleneck in common; they lack the possibility for parallelization and high-throughput set-ups. Indeed, enrichment strategies such as (density gradient) ultracentrifugation require (multiple) lengthy centrifugation steps, are generally limited to six or eight samples and require expensive instrumentation. Size-exclusion chromatography (SEC) often requires a concentration step both prior and after EV enrichment and, in addition, also validation (often by Western blotting) to identify those fractions containing EV material (50). Commercial EV enrichment kits are again expensive and, more importantly, show high variability between suppliers (51,52).

Ultrafiltration (UF) has thus far been mainly used as an intermediate step in EV enrichment strategies to reduce the volume of conditioned medium or liquid biopsy either prior (e.g. density gradient ultracentrifugation) or after EV enrichment (e.g. SEC) (53–55). The main challenges associated with ultrafiltration for EV proteome analysis are the inefficient removal of non-EV proteins, irreversible blocking of the filter membrane and low recovery of the retained particles (56,57). However, it also offers inherent advantages, such as the limited cost, reduced sample preparation time and the potential of analyzing multiple samples simultaneously using standard lab equipment. In addition, UF has little bias towards EV subpopulations (58), in contrast to (for instance) immuno-capture.

Here, we propose an ultrafiltration strategy for the isolation and purification of EVs from conditioned medium and bovine serum, which we termed filter-aided extracellular vesicle enrichment or FAEVEr. In short, pre-cleared conditioned medium or serum is loaded on 300 kDa MWCO filters and centrifuged briefly at a moderate speed during which EVs are retained on the filter, whereas the bulk of globular (non-EV) proteins do not. Remaining filter-retained non-EV proteins are removed through subsequent purification steps that include TWEEN-20-containing wash buffers. The retained (purified) EVs are then efficiently lysed on the filter in SDS-containing buffer and the EV lysates are conveniently collected by centrifugation and processed for proteomics applications, such as Western blotting and LC-MS/MS analysis. Recombinant EV (rEV) material isolated from HEK293T conditioned medium was used for testing and optimizing our strategy for comparison to ultracentrifugation, the most widely implemented EV enrichment strategy. We demonstrate that the EV purity is considerably improved when including TWEEN- 20 in the wash steps without jeopardizing EV integrity, even at high concentrations of the detergent. We then successfully applied FAEVEr to complex biofluids (bovine serum) and investigated the effect on the EV protein cargo when using serum-depleted medium on MCF7 cell cultures.

## B. Material and methods

### 1. Cell culture

All cell culture experiments were performed using standard protocol and lab equipment.

#### a. rEV production

A complete protocol for the transformation and production of recombinant extracellular vesicles (rEV) material is described in (49,59). In short, approximately 4 million HEK293T cells of low passage number (< 10) were seeded in a T75 falcon with Dulbecco’s Modified Eagle Medium (DMEM, Gibco) containing 10% standard FBS and incubated for 48 h at 37 °C in 5% CO2. The HEK293T cells were transfected by adding a mixture of polyethyleneimine (PEI) and the DNA bait construct (12x, pMET7- GAG-eGFP) in DMEM + 2% FBS for 6 h at 37 °C in 5% CO2. The medium was then discarded and replaced with 8 mL fresh DMEM supplemented with 10% EV-depleted FBS (EDS, Thermo A2720801). After 48 h incubation at 37 °C in 5% CO2, cell transfection was evaluated under UV light (excitation at 488 nm and emission at 507 nm) before the conditioned medium (CM) was isolated. The CM was immediately pre-cleared by centrifugation at 1,000 x g for 10 min and filtration of the supernatant using 0.22 µm filters (Millex). The pre-cleared CM was divided over different aliquots and frozen in -80°C until further use. Note that pure rEV extracts are commercially available (SAE0193, Sigma-Aldrich).

#### b. MCF7 cells

MCF7 cells of low passage number (< 10) were cultured using a standard protocol and equipment. The cells were seeded in T75 flasks with DMEM containing 10% standard FBS and incubated at 37 °C in 5% CO2. When the cells reached 70-80% confluence, the medium was discarded and the cells were washed three times with PBS. Experiments were performed in duplicate, adding 8 mL of DMEM supplemented with either 0%, 2%, 5% or 10% EDS (Thermo, A2720801) or OptiMEM supplemented with 0% EDS. After 24 h, the CM containing 0% EDS were isolated, whereas the 2%, 5% and 10% EDS containing CM were isolated after 48 h. The CM was immediately pre-cleared by two centrifugation rounds (500 x g for 5 min, and 10,000 x g for 10 min, room temperature) and filtration using 0.22 µm filters (Millex).

### 2. EV enrichment

#### a. Ultrafiltration using 300 kDa MWCO filters

Pre-cleared CM was diluted 1:1 with PBS with or without supplemented TWEEN-20 and vortexed for 5 – 10 s. The final volume was then transferred to 300 kDa MWCO polyethylene sulfone (PES) filters (Vivaspin®6, Sartorius VS0652) and centrifuged. Centrifugation was done for 10 – 15 min at room temperature either at 6,000 x g for fixed angle rotors (F14x50cy, Sorvall Lynx 4,000, Thermo) or at 4,000 x g for swing-out rotors (e.g. A-4-44, 5804R, Eppendorf). The filter retentate was washed several times with wash buffer (PBS with or without supplemented TWEEN-20) and centrifuged for 5 – 7 min per wash step. After three rounds of washing, the retained particles were either solubilized from the filter for nanoparticle tracking analysis or electron microscopy, or immediately lysed on the filter for maximal recovery of the proteome. For recovery, 500 µL 50 mM HEPES was added on the filter and vortexed for 10 – 15 sec after which the solution was transferred to a fresh tube and vacuum-concentrated (SpeedVac). To immediately collect the EV lysate, 5% SDS in 50 mM TEAB is added and collected after 15 min incubation (37°C, 500 rpm) by centrifugation (5 min at room temperature).

#### b. Ultracentrifugation

Pre-cleared conditioned medium was transferred to polycarbonate thick-wall tubes (Beckman, 343778) and centrifuged for 90 min at 100,000 x g (4°C) using a Optima TLX ultracentrifuge equipped with a TLA-120.2 rotor (Beckman). The supernatant was carefully removed and the pellet washed with PBS with or without TWEEN-20 before a second round of ultracentrifugation for 90 min at 100,000 x g (4°C). Again, the supernatant was carefully removed and the pellets were either solubilized for NTA or EM, or immediately lysed for maximal recovery of the proteome. The EVs were lysed by adding 5% SDS in 50 mM TEAB and the lysate collected after 15 min incubation (37 °C, 500 rpm).

### 3. Proteomics

#### a. Western blot

Samples for Western blot analysis were incubated for 5 min at 95 °C with XT sample buffer (4x) and XT reducing agent (20x) (BioRad). The protein material was separated by SDS-PAGE (4%- 12%) for 80 min at 120 V prior to transfer onto a PVDF membrane (100 V, 30 min). The membrane was incubated overnight at 4 °C with primary antibodies (rabbit monoclonal anti-Calnexin (ab133615) and mouse monoclonal anti-HIV p24 39/5 4A (ab9071); 1:1,000). The next day, the membrane was washed several times before incubation for 1 h at room temperature with secondary antibodies (goat anti-rabbit IRDye 680 CW and goat anti-mouse IRDye 800 CW; 1:10,000). Visualization was done on an Odyssey infrared imager (v3.0.16, LI-COR Biosciences).

#### b. LC-MS/MS sample preparation

The proteome was prepared for LC-MS/MS analysis using S-Trap™ mini columns (ProtiFi, C02- mini-80) using the manufacturer’s protocol. In brief, proteins in lysis buffer (5% SDS 50 mM TEAB, pH 7.4) were reduced and alkylated using 5 mM TCEP (10 min at 55 °C) and 20 mM iodoacetamide (IAA, 10 min at room temperature in the dark), respectively. The sample was acidified to 1.2% phosphoric acid and diluted seven-fold with binding/wash buffer (90% methanol in 100 mM TEAB, pH 7.4). The samples were transferred to the corresponding S-Trap™ columns and each time centrifuged at 4,000 x g for 30 s during sample loading and washing. For protein digestion, 0.5 µg of trypsin (Promega, V5111) was added to 125 µL 50 mM TEAB (pH 7.4) and samples were incubated overnight at 37 °C. The next day, the peptides were recovered by sequential elution by centrifugation (4,000 x g for 30 s at room temperature) using 80 µL 50 mM TEAB (pH 7.4), 80 µL 0.2% formic acid and 80 µL of 50% acetonitrile (ACN), 0.2% formic acid in dH2O. The eluates (app. 365 µL) were transferred to MS vials, vacuum-dried in a SpeedVac and stored at -20 °C. We have submitted all relevant data of our experiments to the EV-TRACK knowledgebase (EV-TRACK ID: EV240045) [34].

#### c. LC-MS/MS analysis

Purified peptides were re-dissolved in loading solvent A (0.1% TFA in water/ACN (98:2, v/v)) and the peptide concentration was determined on a Lunatic instrument (Unchained Lab). For each sample the injection volume was adjusted to inject equal amounts of peptide material (500 ng) for LC-MS/MS analysis on an Ultimate 3000 RSLCnano system in-line connected to a QExactive Exploris (for optimization and validation) or an Orbitrap Fusion Lumos mass spectrometer (Thermo) (MCF7 cells and EVs). QCloud was used to control instrument longitudinal performance during the project [57, 58]. The mass spectrometry proteomics data have been deposited to the ProteomeXchange Consortium via the PRIDE [59] partner repository with the dataset identifiers PXD051938 (comparison FAEVEr with UC), PXD051955 (comparison MCF7 proteome of EVs and cells under starving conditions) and PXD051956 (comparison FBS with EDS).

##### LC-MS/MS analysis using the Fusion Lumos instrument

LC-MS/MS data-independent (DIA) acquisition on the Fusion Lumos was initiated by trapping the peptide material at 20 μl/min for 2 min in loading solvent A on a Pepmap column (300 μm internal diameter (I.D.), 5 μm beads, C18, Thermo). The peptides were separated on a 110 cm µPAC prototype column (Thermo), kept at a constant temperature of 50 °C. Peptides were eluted by a non-linear gradient starting from 2% MS solvent B (0.1% formic acid (FA) in acetonitrile), reaching 26.4% MS solvent B in 82 min and 44% MS solvent B in 90 min and 56% MS solvent B in 100 min starting at a flowrate of 600 nl/min for 5 min, and completing the run at a flow rate of 300 nl/min, followed by a 5-minute wash at 56% MS solvent B and re-equilibration with MS solvent A (0.1% FA in water). The mass spectrometer was operated in data-independent mode. Full-scan MS spectra ranging from 400-900 m/z with a target value of 4E5, a maximum fill time of 50 ms and a resolution of 60,000 were followed by 50 quadrupole isolations with a precursor isolation width of 10 m/z for HCD fragmentation at an NCE of 34% after filling the trap at a target value of 4E5 for maximum injection time of 54 ms. MS2 spectra were acquired at a resolution of 30,000 at 200 m/z in the Orbitrap analyser without multiplexing. The isolation intervals ranged from 400 – 900 m/z, without overlap.

##### LC-MS/MS analysis using the Orbitrap Exploris 240 instrument

The peptide material was injected for LC-MS/MS analysis on a Vanquish™ Neo UHPLC System in-line connected to an Orbitrap Exploris 240 mass spectrometer (Thermo). Injection was performed in trap-and-Elute workflow in combined Control mode (maximum flow of 60 µl/min and maximum pressure of 800 bar) in weak wash solvent (WW, 0.1% trifluoroacetic acid in water/acetonitrile (ACN) (99.5:0.5, v/v)) on a 5 mm trapping column (Thermo scientific, 300 μm internal diameter (I.D.), 5 μm beads). The peptides were separated on a 250 mm Aurora Ultimate, 1.7µm C18, 75 µm inner diameter (Ionopticks) kept at a constant temperature of 45 °C. Peptides were eluted by a gradient starting at 0.5% MS strong wash solvent (SW) (0.1% FA in acetonitrile) reaching 26% MS SW in 30 min, 44% MS SW in 38 min, 56% MS SW in 40 min followed by 5-minute wash at 56% MS SW and column equilibration in Pressure Control mode (separation column: fast equilibration, maximum pressure of 1500 bar, equilibration factor=2; Trap column: fast wash and equilibration, wash factor=100) with MS WW . The flow rate was set to 300 nl/min. The mass spectrometer was operated in data-independent mode, automatically switching between MS and MS/MS acquisition. Full-scan MS spectra ranging from 400-900 m/z with a normalized target value of 300%, a maximum fill time of 25 ms and a resolution at of 60,000 were followed by 30 quadrupole isolations with a precursor isolation width of 10 m/z for HCD fragmentation at an NCE of 30% after filling the trap at a normalized target value of 2000% for maximum injection time of 45 ms. MS2 spectra were acquired at a resolution of 15,000 with a scan range of 200- 1800 m/z in the Orbitrap analyser without multiplexing. The isolation intervals were set from 400 – 900 m/z with a width of 10 m/z using window placement optimization. EASY-IC^TM^ was used in the start of the run as Internal Mass Calibration

#### d. Database searching and data analysis

Data-independent acquisition (DIA) spectra were searched with the DIA-NN software (v1.8.2b) [60] in library-free mode against the combination of two protein databases downloaded from Swiss-Prot; the complete human protein sequence database (January 2021, 20,394 sequences) supplemented with bovine serum protein sequences (March 2021, 618 sequences). For searches that included rEV material, we included the Gag-eGFP protein sequence as well. The mass accuracy was set to 10 ppm and 20 ppm for MS1 and MS2, respectively, with a precursor FDR of 0.01. Enzyme specificity was set to trypsin/P with a maximum of two missed cleavages. Variable modifications were set to oxidation of methionine residues (to sulfoxides) and acetylation of protein N-termini. Carbamidomethylation of cysteines was set as a fixed modification. The peptide length range was set to 7-30 residues with a precursor charge state between 1 and 4 and an m/z range between 400-900 and 200-1,800 for the precursor and fragment ions, respectively. Cross-run normalization was set to RT dependent with the quantification strategy set to high accuracy and the neural network classifier to single-pass mode. The result file was further processed in KNIME (v4.3.3) by removing non-proteotypic peptides and identifications with a protein q-value above 0.01. Peptide quantifications were aggregated to protein group quantifications using the median of the corresponding normalized LFQ values. Further data analysis was performed with Perseus (version 1.6.14.0), GraphPad Prism (v9.4.1) and RStudio (2023.12.1).

For the analysis of the EV and cellular proteomes across different experimental conditions, proteins that were significantly differentially abundant were identified by multiple sample testing (ANOVA) and selected for hierarchical clustering (Euclidean distance with averaged linkage) after Z-scoring the individual proteins over the different conditions. Row clusters were defined manually and used for gene ontology (GO) enrichment against the corresponding initial database and included biological process (GOBP), molecular function (GOMF), cellular component (GOCC) annotations and KEGG pathways. GO term enrichment was done using the online WebGestalt tool for Gene ontology enrichment (GOA) (60).

### 4. EV characterization

#### a. NTA analysis

Recuperated EV particles were diluted to 1 mL using PBS (pH 7), injected in a calibrated Zetaview and analyzed with the corresponding software package (version 8.05.16 SP2). The temperature was maintained at 23°C and the analysis was done at a sensitivity of 70 and a shutter of 100 for 3 cycles at 60 fpm. The results were plotted and analyzed in GraphPad Prism (v10.2.2)

#### b. Scanning electron microscopy

Filters containing EVs were placed in 2 ml Eppendorf tubes and fixed with 4% paraformaldehyde and 2% glutaraldehyde in 0.1 M Sodium cacodylate buffer at room temperature. After one hour, the fixative was replaced and filters were washed 3 times for 5 min with dH2O. After dehydration in 30%, 50%, 70%, 95% and 2x 100% ethanol for 30 min each, the filters were critical point dried (EM CPD300, Leica) and imaged on a Crossbeam 540 SEM (Zeiss) at 1kV.

#### c. Transmission electron microscopy

Aliquots (5 µl) of the EV solutions were blotted for 1 min on formvar- and carbon-coated Ni maze grids (EMS), which were glow discharged for 40 sec at 15 mA. The grids were then washed five times in droplets of dH2O and stained in a droplet of 1/4 Uranyl Acetate Replacement Stain (EMS)/ dH20 for 1 min. Excess stain was removed with filter paper and the grids were air dried for at least four hours before viewing with the TEM. Imaging was done at 80kV on a JEM1400plus (JEOL).

## C. Result section

### 1. Use of 300 kDa MWCO filters to enrich EVs

Although ultrafiltration is a valid and well-implemented method to concentrate intact EVs, it holds many challenges as a stand-alone EV enrichment strategy. Nevertheless, ultrafiltration also has important advantages such as low cost and fast processing times, and it does not require specific instrumentation compared to, for instance, ultracentrifugation.

Here, we introduce Filter-Aided Extracellular Vesicle Enrichment (FAEVEr), a strategy that uses 300 kDa MWCO polyether sulfone (PES) filters to purify and concentrate intact EVs (Figure 1). In brief, pre-cleared conditioned media or biofluids are transferred onto the 300 kDa MWCO filter membrane and centrifuged at moderate speed (4,000 – 6,000 x g), retaining the EV particles on the filter (retentate) whereas globular non-EV proteins are removed (filtrate). By including three consecutive wash steps, non-EV proteins are increasingly removed, by which the EV retentate is further purified. Eventually, by using a lysis buffer with high percentages (5%) of SDS, EVs undergo efficient lysis and their cargo is conveniently recovered by centrifugation. The EV sample is then further processed for proteomics using S-Trap columns (61). Here, we opted for PES membranes, as filters based on such membranes are hydrophilic with low protein adsorption characteristics. In addition, PES material is resistant against high pH ranges and temperatures and can withstand a wide variety of solvents, buffers and detergents.

**Figure 1:**
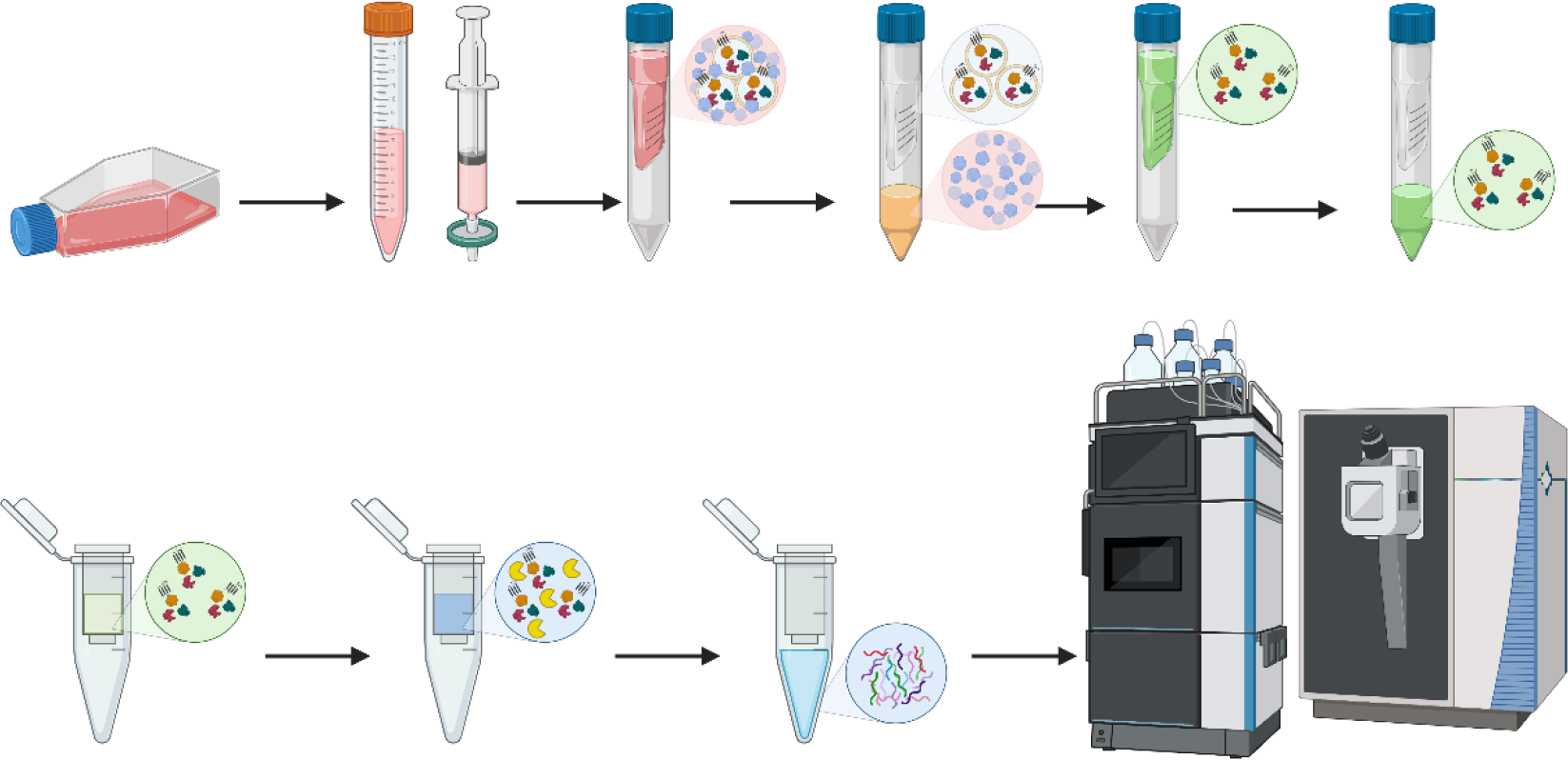
Schematic overview of the FAEVEr workflow. Conditioned media or biofluids are precleared and transferred onto the 300 kDa MWCO PES filter (red). Driven by centrifugation, the globular proteins pass through the filter (orange), whereas the EV particles are quantitatively retained (grey). The retentate is subsequently washed three times using PBS supplemented with TWEEN-20 to further increase the EV purity. Ultimately, intact particles are recovered for qualitative analysis, or lysed (green) immediately on the filter. The lysate is recovered by centrifugation and prepared for proteomic applications (e.g. LC-MS/MS analysis or Western blotting). Figure created in Biorender.

Recombinant EV (rEV) material was used for the initial optimization of the 300 kDa MWCO ultrafiltration strategy as it has similar biochemical and biophysical characteristics as sample EVs (48). The rEV material is derived from the conditioned medium of HEK293T cells, cultured in 10% EV-depleted serum, that overexpress the Gag-eGFP fusion protein (49). The Gag subunit multimerizes at the plasma or endosomal membrane and initiates membrane curving, resulting in the formation of microvesicles and intraluminal vesicles, respectively, commonly referred to as virus-like particles (59,62) (Error! Reference source not found.). As a result, secreted rEV material is overrepresented in the conditioned medium and contains high levels of luminal Gag-eGFP, enabling the evaluation of EV integrity during the consecutive FAEVEr steps.

We confirmed that rEV particles are retained on the 300 kDa MWCO mesh using scanning electron microscopy (SEM) (Figure 2A and Error! Reference source not found.), indicating that ultrafiltration is an efficient EV enrichment strategy. Indeed, after completing the FAEVEr protocol, rEV particles were recovered from the filter for nanoparticle tracking analysis (NTA) (Figure 2B) and transmission electron microscopy (TEM) characterization (Figure 2C). However, in line with Vergauwen *et al*. (56), we noticed that the quantitative recuperation of particles from the filter PES membrane is variable and that not all particles are recovered. When the remaining rEV fraction on the filter was lysed and analyzed on Western blot, we observed that a substantial share of the rEV material fails to be solubilized and remains on the filter (Figure 2D).

**Figure 2:**
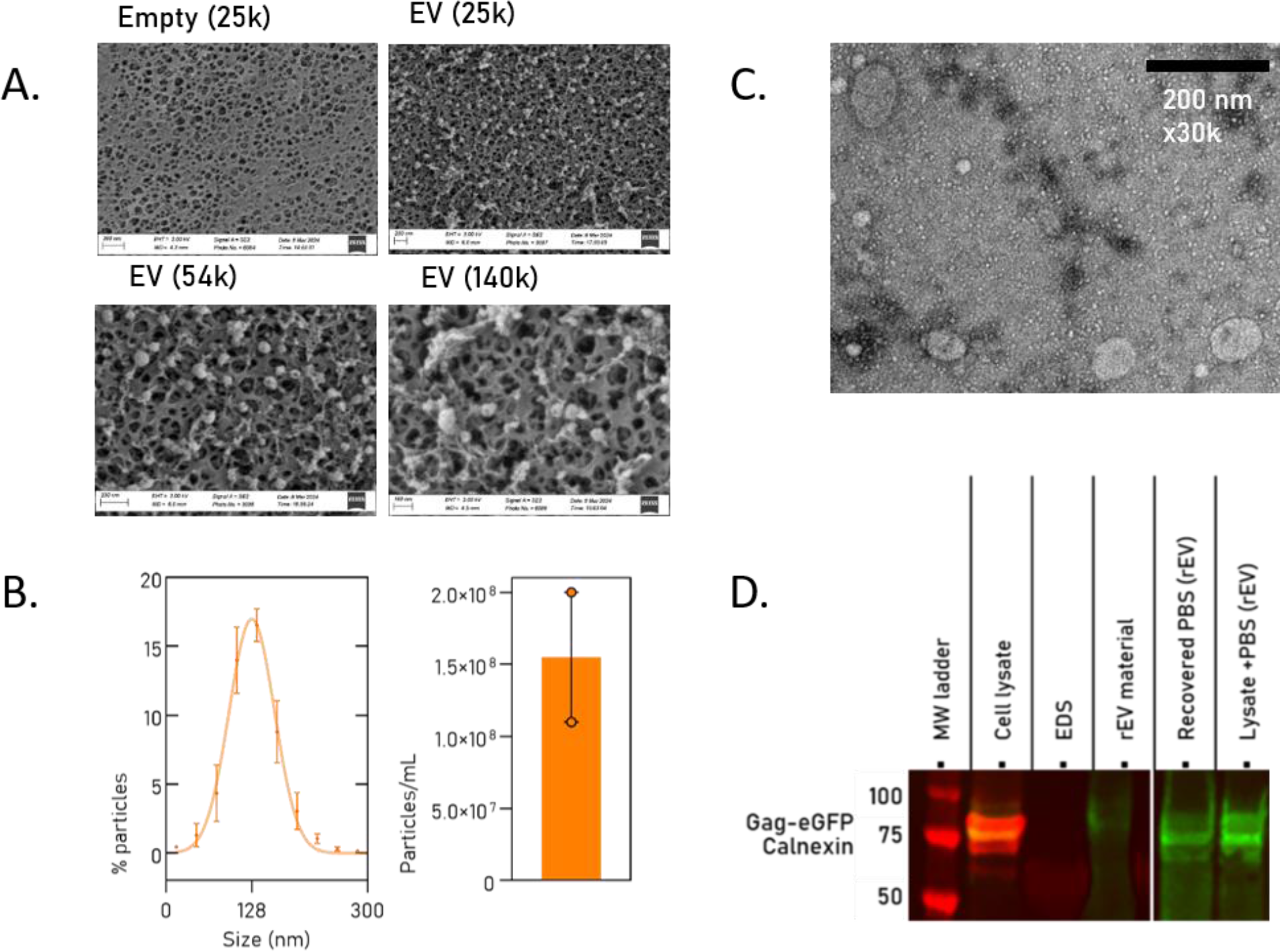
Qualitative analysis of FAEVEr retained particles. A) Scanning electron microscope of an empty filter and after rEV enrichment. B) EV size distribution and count, and transmission electron microscopy (C) of recovered rEV particles from the filter. D) Western blot reveals incomplete recuperation of rEV particles from the filter. EDS: EV-depleted serum (negative control), rEV material: commercial rEV particles (positive control), PBS: phosphate-buffered saline washed rEV particles.

Concluding this part, we found that FAEVEr is an efficient approach for isolating EV particles from conditioned medium for downstream proteome analysis and that we can successfully recuperate intact particles from the filter (56). By immediately lysing the filter-retained EV material using high percentages of SDS for proteomic applications, we ensure EV lysis, comprehensive solubilization of (membrane) proteins (63) and maximal protein recovery. Note that SDS is irreconcilable with LC-MS/MS analysis and we therefore introduced S-Trap columns in our workflow (61,64). The S-Trap protocol is easy, fast and reproducible and has been extensively compared to other methods in different studies (65,66).

### 2. Comparison of FAEVEr to ultracentrifugation

Using EV proteome analysis, we set out to compare our FAEVEr approach with ultracentrifugation (UC), the most used strategy for EV enrichment. rEV material was isolated, pre-cleared and divided over different aliquots before subjecting it to the corresponding EV enrichment strategy in triplicate. In terms of sample preparation, the overall sample preparation time differs noticeably. Ultracentrifugation required at least two rounds of centrifugation, totaling to 3 h of centrifugation time, whereas FAEVEr was finished within the hour, including centrifugation steps and hands-on time.

For an in-depth analysis of the proteome results, we examined the occupancy of the LC-MS/MS spectral space, which is a measure of the recorded spectra and allows us to differentiate between recorded spectra of bovine and human origin. Our results indicate that the relative spectral space occupancy by bovine precursors was slightly reduced (from 35% to 30%) using FAEVEr compared to UC (Figure 3A). In addition, approximately 15% more proteins were identified proportionally distributed over the different cellular locations (Figure 3B). The six commonly used EV markers are well represented and identified with similar relative intensities between strategies (Figure 3C) and 85% of the identified proteins by the UC were found by FAEVEr (Figure 3D). In addition, the same trend was observed for the number of unique peptides (13%) and the number of the associated peptide-spectral matches (PSMs), the number of recorded spectra that were translated in a confident peptide identification (14%) (Figure 3E). Importantly, we observed a large decrease in the accompanying coefficient of variance (%CV) for FAEVEr between the replicates as well as fewer missing values, which results in an increased data completeness and therefore higher reproducibility with more accurate protein quantifications (Figure 3F). From this experiment, we can thus conclude that ultrafiltration using 300 kDa MWCO filters allows for EV enrichment and is comparable to UC in terms of EV proteome depth.

**Figure 3:**
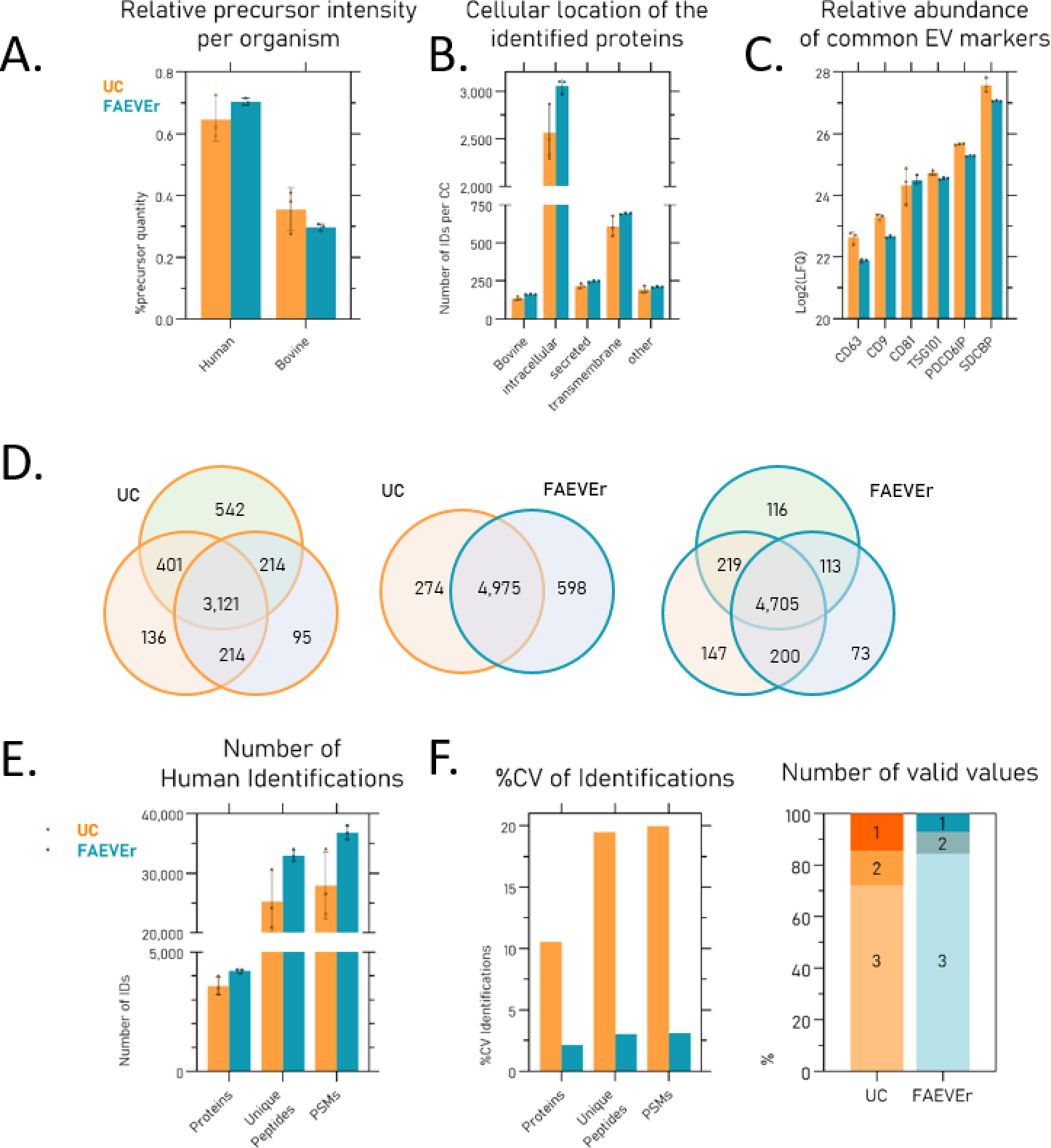
Overview of the comparison between FAEVEr and ultracentrifugation. A) Comparison of the relative abundance of precursor material from bovine or human origin. B) Distribution of the protein identifications across the cellular localizations of the parental cell. C) High relative abundance of six molecular EV markers. D) Overlap of protein identifications between UC and FAEVEr. E) Total number of identified human proteins, identified unique peptides and peptide- spectral matches from these proteins between UC and FAEVEr. F) Decreased relative variance (%CV) and fewer missing values are observed with FAEVEr compared to UC. UC: ultracentrifugation (orange), FAEVEr: Filter-aided EV enrichment (teal), CC: cellular compartment, LFQ: label-free quantification intensity, PSM: peptide-spectral match, %CV: coefficient of variance.

### 3. TWEEN-20 improves the removal of non-EV proteins

One major pitfall of ultrafiltration is the inefficient removal of non-EV proteins, which has a detrimental effect on the numbers of identified and quantified EV proteins following LC-MS/MS analysis. Therefore, we explored the use of mild detergents to minimize the interaction between proteins and the filter membrane. However, as EVs are delineated with a lipid bilayer and therefore susceptible to detergent-mediated lysis, caution should be taken with the choice and concentration of the detergent. A study by Osteikoetxea *et al*. explored the minimal concentration of different detergents required for efficient lysis of EV particles in solution (67) and found that this concentration greatly depends not only on the detergent used, but also on the EV subpopulation (exosomes, microvesicles and apoptotic bodies). In particular, the smallest population (exosomes) demonstrated increased resilience against detergents compared to larger vesicles (microvesicles and apoptotic bodies) due to an increase in liquid order because of an accumulation of cholesterol and sphingolipids during their formation (68–70). In their work, Osteikoetxa *et al.* demonstrate that EVs are generally efficiently lysed even at low percentages of SDS, Triton X-100 or sodium deoxycholate. On the contrary, an exceptional high tolerance is observed towards TWEEN-20, a rather mild detergent that is commonly used in other biochemical assays such as Western blotting to reduce unwanted adsorption of protein material to membranes (71) and thereby decreases irreversible or total membrane fouling (72,73). TWEEN-20 was explored here as an additive to improve the efficiency of the washing steps and thus removal of non-EV proteins during the enrichment of EVs whilst keeping EVs intact. The efficiency of washes with TWEEN-20-containing buffers was tested by Coomassie staining of the different filtrate fractions of conditioned medium after completing the FAEVEr protocol using 0%, 0.1%, 0.5%, 1%, 2% and 5% TWEEN-20 in the wash buffer. We observed that PBS-only (the 0% TWEEN-20) condition is not capable of removing the bulk of bovine serum albumin (BSA; 65 kDa), whereas BSA-bands are more intense in the TWEEN-20 supplemented wash fractions. In addition, upon recovery of the retained material by lysis in 5% SDS, we observed an intense band for the PSB-only condition, which was absent in the TWEEN-20 supplemented ones (Error! Reference source not found.). This suggested that TWEEN-20 is indeed capable of removing the bulk of (contaminating) bovine serum material from the filter membrane and therefore should result in an EV proteome fraction that is more pure.

Next, we validated this beneficial effect of TWEEN-20 in wash buffers using rEV material, which also allows us to validate the integrity of the particles as they contain the Gag-eGFP protein on the luminal side. We thus argued that, when the rEV particles remain intact, no luminal Gag- eGFP should be present in the filtrates. In short, pre-cleared conditioned medium containing rEV material was filtered (300 kDa MWCO) and washed several times with up to 5% TWEEN-20 in PBS. After each centrifugation step, the filtrate was collected and the presence of luminal Gag- eGFP validated on Western blot, shown in Figure 4A. The almost complete absence of luminal Gag-eGFP in the flow-through and washes (1–3) does not only indicate that intact rEV particles are retained on the filter, but also that they maintain their integrity in concentrations up to 5% TWEEN-20, as validated by TEM, SEM and NTA (Figure 4B). In addition, it also indicates that the proteome of retained EVs is efficiently recovered from the filter for downstream proteomic applications using a lysis buffer containing 5% SDS.

**Figure 4:**
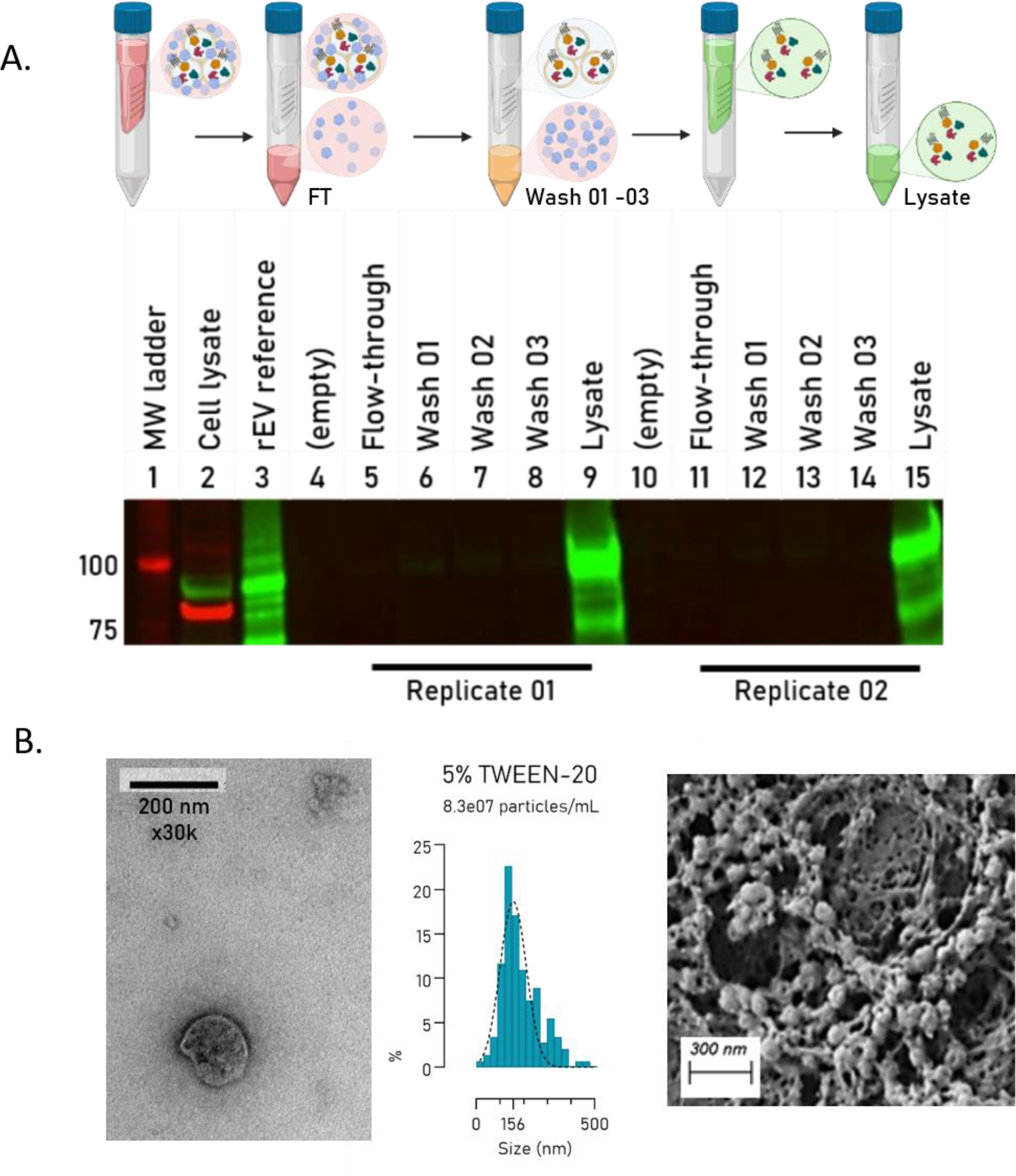
The integrity of rEV particles remains even when washing with buffers supplemented with 5% TWEEN-20. A) Western blot of the different fractions during FAEVEr showing that no (or just marginal) luminal Gag-eGFP is observed in the flow-through and wash steps, implying that EVs remain intact. In addition, a strong signal is observed in the lysate fractions, indicating that we can efficiently recover the rEV proteome. B) Qualitative analysis by TEM, NTA and SEM of recovered rEV particles washed in PBS supplemented with 5% TWEEN-20.

We further extended our comparison between FAEVEr and UC to evaluate the effect of TWEEN- 20 on the purity of the final rEV proteome. We included different percentages of TWEEN-20 in the washing buffer of both FAEVEr and UC and analyzed the corresponding proteomes by LC-MS/MS. We initiated the analysis by comparing the spectral space distribution between human and bovine precursors. Here, we did not only observe a remarkable decrease in bovine precursor intensities using FAEVEr even at the lowest (0.1%) TWEEN-20 concentration, but also that increasing the percentage further improved this (Figure 5A). The effect of supplementing TWEEN-20 is summarized in a principle component analysis (PCA, Figure 5B) which illustrates that supplementing TWEEN-20 has the most drastic change in combination with FAEVEr.

**Figure 5:**
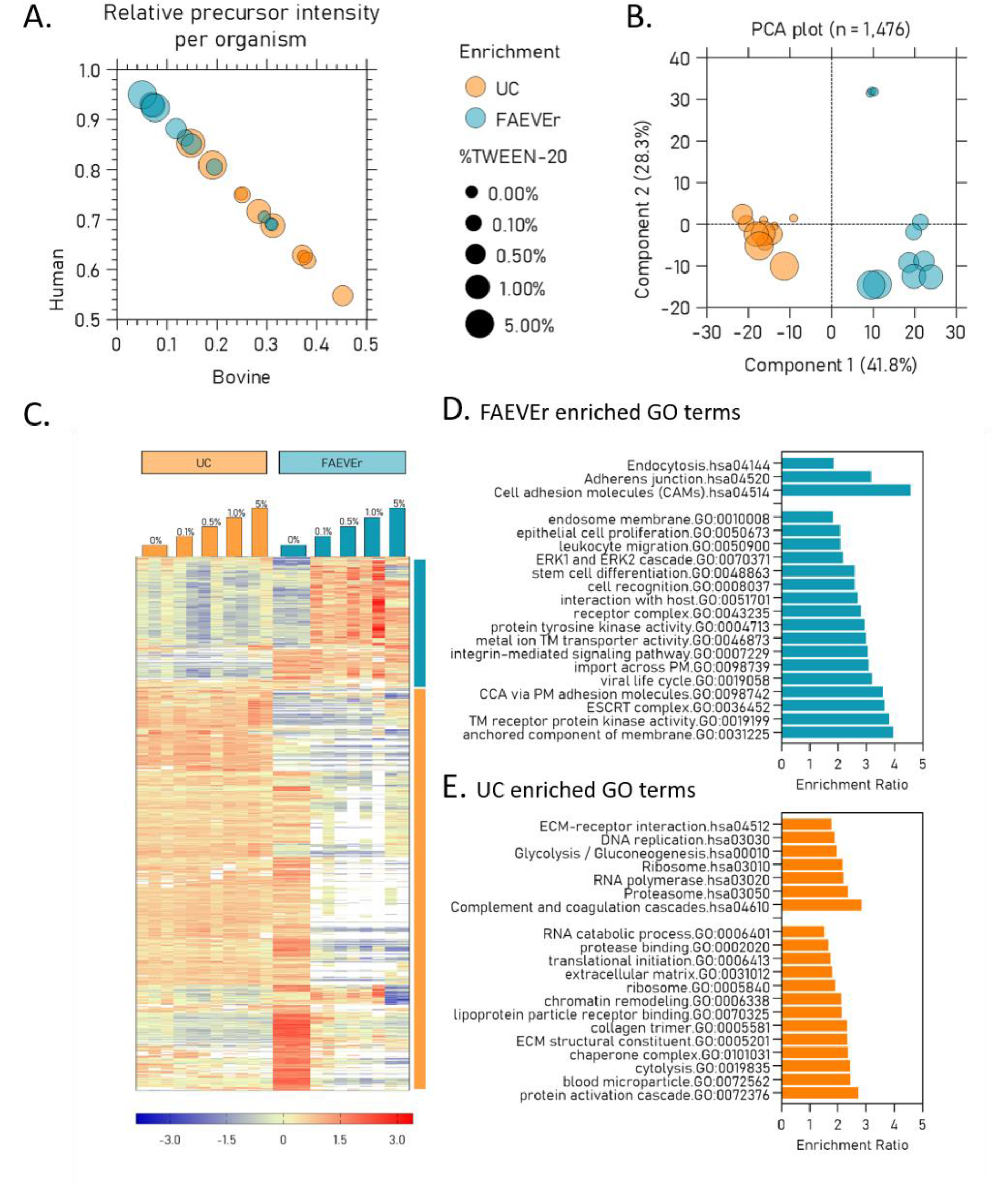
Overview of the effect of TWEEN-20 at different percentages on EV enrichment by UC and FAEVEr. A) Relative share of bovine and human precursor abundance as a measure of serum contamination in function of the enrichment strategy and the percentage of TWEEN-20 in the wash buffer. B) PCA plot of the shared protein intensities between the different experimental setups reveals three major groups. C) Heatmap of the significantly differential abundant proteins (multiple sample test, p-value < 0.05) per experimental setup with the accompanying GO analysis for FAEVEr enriched (D) and UC enriched (E) proteins. UC: ultracentrifugation (orange), FAEVEr: filter-aided EV enrichment (teal), PCA: principal component analysis, GO: Gene ontology, TM: transmembrane, CCA: cell-cell adhesion, ECM: extracellular matrix.

Indeed, by combining FAEVER with 5% TWEEN-20, we identified over 3,700 proteins with nearly 95% of all PSMs to be of human origin. For UC supplemented with 5% TWEEN-20 in the wash buffer, we also observed an improvement in terms of decreased serum contamination, although not to the same extent, where approximately 80% of the observed precursors were of human origin. Interestingly, we identified over 4,400 proteins in the UC setup, or roughly 15% more proteins in comparison to FAEVEr. We investigated this change in protein abundance further as we expected these extra proteins not all to be serum proteins. We found, by initiating a multiple sample test between the individual experimental setups, (as visualized in the PCA plot) three major groups; UC with or without TWEEN-20, FAEVEr without TWEEN-20 and FAEVEr with TWEEN-20. The setup in which FAEVEr was used without TWEEN-20 appeared to associate with both groups to some extent (Figure 5C). By further examining gene ontology and KEGG terms associated with the corresponding proteins, we found that in FAEVEr supplemented with TWEEN-20, terms including cell-cell adhesion, ESCRT mechanism and (trans)membrane proteins were more abundant (Figure 5D). On the other hand, we noticed that in UC, terms including RNA binding, cell cycle, proteasome and metabolic activities were more prominent (Figure 5E) as well as serum-related terms (e.g. blood microparticle, activation cascade).

In conclusion, our results indicate that FAEVEr in combination with TWEEN-20 in the washing buffer improves the overall EV proteome purity, whereas this effect is less prominent in UC.

### 4. Effect of serum depletion (on the EV proteome)

Fetal bovine serum (FBS) is used as a standard supplement in cell culture media. Unfortunately, it is irreconcilable with EV analysis from conditioned cell media as it contains EVs from bovine origin that interfere with proteome analysis. Therefore, EV depleted serum (EDS) alternatives are used. Consequently, researchers often expand cells for several days in media containing EDS or even refrain from adding serum. To this day, it remains a controversial topic (74) with some researchers advocating against the use of serum because it remains a potential source of xeno-contamination. Others deem serum as an essential supplement for normal cell growth with serum depletion stressing cells due to starvation (75,76), which potentially biases the proteome of the cell culture-derived EVs. Benefiting from the EV-TRACE repository, we noticed that between 2018 and 2023 approximately 56% of the EV experiments that were conducted on conditioned medium of human cell lines were performed using EDS, whereas 30 – 35% of the EV experiments was performed using serum-free conditions.

Following the MISEV2018 (7) guidelines, we investigated whether the used EDS is indeed devoid of EV material and addressed this using FAEVEr comparing EDS with FBS. In addition, we examined uses of FBS or EDS had an effect on the cellular and the EV proteome. For these experiments, we made use of MCF7 breast cancer cells.

As the high protein concentrations and the accompanying viscosity of serum samples may result in low filtration speed and irreversible blocking of the filter membrane, we diluted the sample three times prior to filtration. After completing the FAEVEr protocol, the proteomes of the FAEVEr filter-retained fraction from commercial FBS and EDS were compared by LC-MS/MS. We validated that EV-specific molecular markers were absent in EDS as well as proteins associated with EV biogenesis. In contrast, these proteins were found abundantly in the FBS retained fraction (Error! Reference source not found.).

In conclusion, using FAEVEr, we here provide evidence at the LC-MS/MS level that EDS is indeed devoid of EV material and that the FAEVEr strategy is capable of enriching EV particles from complex biological fluids such as (bovine) serum.

The protein concentration of serum is reduced by a factor six because of the EV depletion process (Error! Reference source not found.A). Indeed, FBS is diluted several times to prevent unintended protein aggregation and pelleting by ultracentrifugation to remove EVs due to the high viscosity of undiluted FBS (73). We decided to explore the effect on the MCF7 cellular proteome once the medium was exchanged from 10% FBS to 10% EDS for 3 days. Besides no change in cell morphology by visual inspection or a significant change in cell count or cell death (Error! Reference source not found.B), the LC-MS/MS analysis indicated that only 10 out of more than 7,000 quantified proteins appeared to be significantly differentially regulated between the two conditions (Error! Reference source not found.C & 5D). We conclude that shifting from FBS to EDS close to no influence on the viability, growth or proteome of MCF7 cells in culture.

We continued investigating the use of FAEVEr by exploring the effect of starvation on the EV proteome by isolating EVs from the conditioned medium of MCF7 cells grown in DMEM supplemented with 0%, 2%, 5% or 10% EDS (D0, D2, D5 and D10, respectively). The FAEVEr retained EVs were purified with wash buffer containing 5% TWEEN-20 and either recovered for NTA (Figure 6A) or lysed for LC-MS/MS analysis. The LC-MS/MS generated data immediately hinted towards an important difference by protein content and quantity between serum-deprived and EDS containing conditions as indicated in the number of identifications (Figure 6B) and the PCA plot (Figure 6C). Over 5,191 proteins were identified in total, from which 4,887 were human and 304 were bovine. From the human proteins, 3,610 were found in duplicate in at least one experimental set-up and were included for quantitative comparison of the EV proteomes. Interestingly, we found that only 959 proteins were identified across all samples and that over 1,100 proteins were uniquely identified in D0. In addition, we found that the relative abundance of transmembrane proteins gradually increased with increasing levels of EDS, whereas levels of intracellular proteins decreased (Figure 6D). EV markers were abundantly picked-up across the individual samples (Figure 6E). Proteins were then quantitatively compared (multiple sample testing) and hierarchical clustered (Figure 6F) which lead to the conclusion that 333 proteins were significantly differential abundant over two main clusters, between the serum depleted and serum containing conditions. These two clusters were used for gene ontology analysis (GOA) using the Webgestalt v2017 online tool (60). Note that the D0 unique protein list was joined with the proteins that were significantly more abundant in the serum-depleted condition. This GOA analysis revealed that EVs isolated from MCF7 cells cultured in serum-depleted conditions for only 24 h are enriched in proteins involved in metabolism, RNA and DNA binding and autophagy, including organelle fission. Indeed, serum deprived cells are known to rapidly adjust the cellular proteome (Error! Reference source not found.), cells are known to upregulate autophagy related (ATG) proteins to replenish essential building blocks and release non-ESCRT dependent exosomes through the formation of autophagosomes (77), which can fuse with endosomes to form amphisomes, similar to MVBs. In addition, the ATG5-ATG16L complex (78,79) influences the EV secretion through dissociation of V-ATPases, thereby preventing MVB acidification and its subsequent lysosomal degradation (80,81). Ultimately, this promotes MVB fusion with the PM and secretion of the ILVs as exosomes. We found that multiple key ATG proteins and contributors of the phagosome biogenesis are high abundant in the EVs isolated from the serum-starved MCF7 cells yet nearly completely absent in the EVs isolated from the serum-containing conditions (Figure 6G & 6H). These findings are in contrast to the serum-containing conditions, where GO terms associated with (trans)membrane proteins (including transmembrane transport, cell-cell adhesion, integrins, vesicle-mediated transport) are enriched as well as proteins associated with the ESCRT complex.

**Figure 6:**
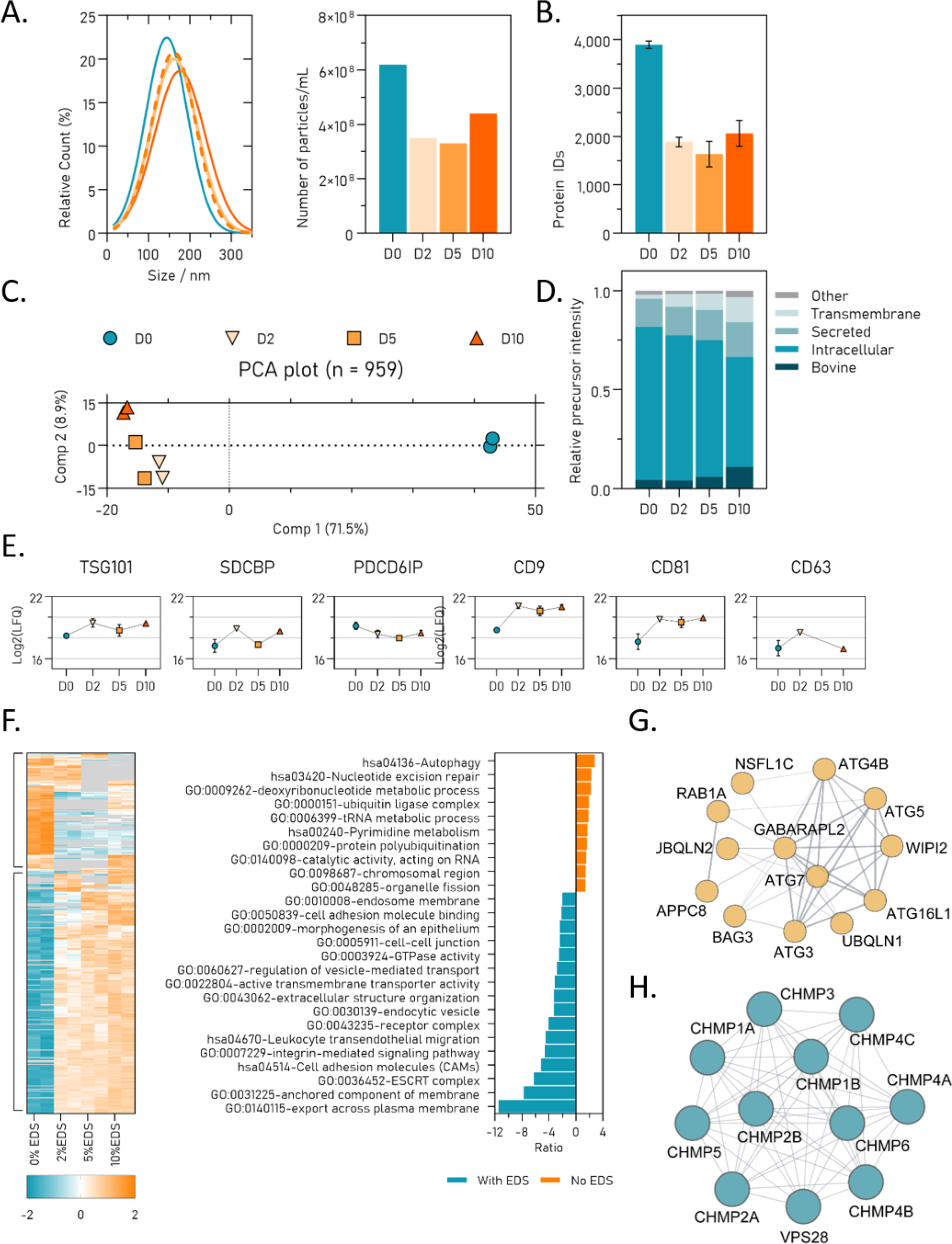
Comparison between the EV proteomes isolated from MCF7 cells cultured in different percentages of EV-depleted FBS (EDS) reveals drastic changes in protein heterogeneity and abundances. A) NTA analysis of recovered EV particles. B) Number of protein identifications between the conditions. C) PCA plot indicating large variation between the relative abundances of proteins identified in serum-starved (D0) and serum-supplemented conditions (D2, D5 and D10). D) Plot of the localization of the identified proteins. E) Profile plot of six commonly used EV markers. F) Heatmap of the significantly differential abundant proteins between the different conditions and the associated enriched GO terms for serum-starved (orange) and serum-supplemented (teal). Detail of ATG related proteins in the serum-starved condition (G) and the ESCRT associated proteins in the serum-supplemented conditions (H). EDS: EV-depleted FBS; D0, D2, D5 & D10: DMEM supplemented with 0%, 2%, 5% and 10% EDS; NTA: nanoparticle tracking analysis; PCA: principal component analysis; GO: Gene ontology.

## F. Conclusion and discussion

Despite the potential of extracellular vesicles (EVs) for clinical applications such as being a source of circulating biomarkers or therapeutic shuttles, their efficient isolation and purification from complex biofluids in a cost and time efficient manner remains a major bottleneck. Here, we propose filter-aided extracellular vesicle enrichment (FAEVEr) as an EV enrichment strategy that uses 300 kDa MWCO filtration. By combining the filtration steps with wash buffers that contain up to 5% TWEEN-20, we were able to improve the EV proteome purity towards LC-MS/MS compatible levels, even in the presence of 10% EV-depleted FBS (EDS). Recombinant EVs (rEVs, (48,49)) were used to initially establish and optimize our strategy. We made use of the overexpression of a luminal Gag-eGFP construct to evaluate the rEV integrity during the enrichment and purification and concluded that during FAEVEr the rEV particles indeed remain intact. In addition, we concluded that the EVs can be recovered from the filter membrane for further evaluation using common tools for EV characterization such as nanoparticle tracking analysis (NTA) and scanning and transmission electron microscopy (SEM and TEM). In addition, the rEV proteome was quantitatively recovered by immediately lysing the filter-retained EVs for LC-MS/MS analysis and we identified over 5,000 human HEK293T proteins from 1 mL conditioned medium. Amongst these proteins we found the different EV markers next to multiple (trans)membrane proteins and proteins associated with EV biogenesis, which underlines the potential of FAEVEr for EV proteomics.

We benchmarked our approach against ultracentrifugation (UC), a widespread implemented method for EV enrichment, and showed that over 95% of the protein identifications overlap. Furthermore, we found an increase in EV purity following FAEVEr compared to ultracentrifugation as demonstrated by a reduced fraction of the LC-MS/MS spectral space taken by bovine precursors, thereby limiting the competitive effect towards the identification of low abundant (EV) proteins. In addition, we observed a drastic decrease in the relative variance (%CV) between the number of identifications on both the protein, peptide and peptide-spectral match level between UC and FAEVEr. This also indicates a higher reproducibility between replicates, which in turn leads to improved data completeness (fewer missing values) and, consequently, improved quantifications. We hypothesize that this improvement is due to the absence of intermediate pipetting steps that remove the supernatant after ultracentrifugation, which potentially leads to incomplete removal or unintended perturbations of the EV pellet. In FAEVEr however, we create a distinct separation between the EV depleted filtrate and EV enriched retentate, thereby omitting unnecessary pipetting steps, as the filtrate is simply decanted.

We continued to improve the EV purity by implementing TWEEN-20 in wash buffers as this detergent was previously shown not to lyse EVs, even in relatively high concentrations (67), and it is an established supplement to reduce non-specific protein-membrane interactions, for example in Western blotting. We provide insights in the gradual loss of bovine serum proteins and secreted proteins, to which we refer here as non-EV proteins. This improved removal of unwanted proteins is of high importance for LC-MS/MS based bottom-up proteomics, as precursors from such high abundant proteins (e.g. albumin) dominate the spectral space, outcompeting relevant EV precursors. Surprisingly, we observed that supplementing TWEEN-20 to the wash step of UC did not have the same profound effect compared to FAEVEr. Indeed, a number of proteins associated with (amongst others) transcription and translation, proteasomal degradation and metabolic activities were more prominent in all ultracentrifugation samples as well as in the FAEVEr setup without TWEEN-20 supplemented to the wash buffers. In contrast, the TWEEN-20 washed EV proteomes following the FAEVEr protocol were enriched in (trans)membrane proteins and in proteins from the ESCRT machinery. We hypothesize that during cell culture inevitable cell lysis occurs during which cytosolic proteins are released and co-purified during ultracentrifugation. These insights raise the question whether or not such proteins are effectively encapsulated in EVs or if they form non-specific protein interactions with the EV membrane. However, additional studies are required to confirm this.

In EV research, many experiments exclude serum supplements in cell culture media to repulse xeno-proteins and omit the suppressive competitive effect during LC-MS/MS analysis and the potential biological influence on the EV proteomics outcome. We first investigated on the difference between regular FBS and EV-depleted FBS (EDS) on the LC-MS/MS level, from which we concluded that EV markers and biogenesis related proteins are virtually absent in EDS, implying that we do not contaminate our EV preparation unintendedly with bovine EV particles.

We further elaborate that the depletion of serum during cell culture, even for a limited period of time (e.g. 24 h), has a dramatic effect on the EV proteome of MCF7 cells. In line with Leidal *et al.* (77) we observed an increase in protein heterogeneity associated with RNA binding, biosynthesis, cell cycle and autophagy, pointing towards an alternative biogenesis of EV particles through the formation of amphisomes. These findings are underlined by the unique presence of several ATG proteins in the EV preparation of serum-starved MCF7 cells. In contrast to this, the EV proteome from MCF7 cells supplemented with EV-depleted FBS, contained higher levels of proteins associated with transmembrane transport, cell-cell adhesion and the ESCRT-machinery were more abundant. We feel that the choice to add or exclude EDS during cell culture for EV research is an important consideration when testing a preconceived hypothesis as it may bias the interpretation of the EV (proteomics) data (82,83).

Our data shows that filter-aided EV enrichment, FAEVEr, using 300 kDa MWCO PES filters is an efficient strategy to isolate EVs from complex medium and that, by including TWEEN-20, we further improved the overall purity of the EV proteome without affecting the EV integrity, even at relatively high TWEEN-20 concentrations. Furthermore, FAEVEr is conducted in a relatively short timeframe and it offers the possibility of parallelization using standard lab equipment. In comparison, ultracentrifugation is limited to six or ten samples and requires at least two rounds of centrifugation.

## G. Supplemental figures legends

Supplementary Figure 1: Schematic overview of the expression of GAG-eGFP in HEK293T cells. Top: HEK293T cells are transfected with a Gag-eGFP construct, encoding for a Gag-eGFP fusion protein that multimerizes at the endosomal and plasma membrane, inducing the formation of virus-like particles packed with eGFP. Bottom: brightfield and eGFP pictures of HEK293T cells after transfection, prior to the isolation of conditioned medium.

Supplementary Figure 2: Scanning electron microscopy of 300 kDa MWCO filters used in FAEVEr. Blank: empty filter membrane with clear visualization of the 300 kDa MWCO filter membrane. Synthetic particles: homogeneous synthetic particles used for NTA calibration were retained on the filter after completing the FAEVEr protocol and serve as a positive control. rEV: retained rEV particles after completing the FAEVEr protocol with a distinct delineation of the EV particles against the 300 kDa MWCO membrane.

Supplementary Figure 3: SDS-PAGE with Coomassie staining indicates quantitative removal of serum proteins with FAEVEr when TWEEN-20 is supplemented to the filter, indicating decreased membrane fouling. In the PBS condition (no TWEEN-20) a faint band is observed at 65 kDa, indicating that only a fraction of the bovine albumin is removed by the wash steps. This stands in contrast to washing with buffers that are supplemented with of TWEEN-20, where more intense bands are observed in the washes. Additionally, the lysate of PBS contains observable more albumin material, further indicating that PBS alone fails to remove the bulk of contaminants contrary to TWEEN-20 were the lysate does not contain such strong signal.

Supplementary Figure 4: Comparison of the identified proteins in standard fetal bovine serum (FBS) and EV-depleted FBS (EDS). A) After completing FAEVEr and subjecting the lysate to LC- MS/MS analysis, we observed a high number of identifications in FBS but not in EDS, indicating a higher protein heterogeneity due to the presence of EV particles. B) EV markers or biogenesis related proteins are virtually absent in EDS, yet highly abundant in FBS.

Supplementary Figure 5: Culturing MCF7 cells in 10% standard fetal bovine serum (FBS) or 10% EV-depleted FBS (EDS) for three days, has no huge effect on the MCF7 cellular proteome. A) Due to the EV depletion protocol, EDS contains approximately six times less protein material. B) No significant changes in cell viability or cell count were observed. C) In both conditions, we identified over 7,000 proteins from which only 10 were significantly differentially regulated (D).

Supplementary Figure 6: The cellular proteome of MCF7 cells cultured at different percentages of EV-depleted FBS (EDS) changes between serum-starved (0% EDS for 24 h) and optimal conditions (10% EDS for 48 h). A) Heatmap depicting the differentially regulated proteins between the different conditions from which we further investigated two major clusters to evaluate the contrast between 0% EDS and 10% EDS. Gene ontology (GO) analysis of the proteins up-regulated in the serum-starved conditions (B) and optimal conditions (C).

## H. List of abbreviations

DDA: Data dependent acquisition
DIA: Data independent acquisition
EC: extracellular
EDS: EV-depleted FBS
ESCRT: endosomal sorting complexes required for transport
EV: extracellular vesicle
FAEVEr: Filter-aided extracellular vesicle enrichment
FBS: fetal bovine serum
GO: Gene ontology
ILV: inter-luminal vesicles
LC-MS/MS: Liquid chromatography coupled tandem mass spectrometry
LFQ: label-free quantification
MVB: multivesicular body
MWCO: molecular weight cut-off
NTA: nanoparticle tracking analysis
PCA: principal component analysis
PES: Polyethersulfone
PM: plasma membrane
PSM: peptide-spectral match
rEV: recombinant EV
SDS: sodium dodecyl sulphate
SEC: size-exclusion chromatography
SEM: scanning electron microscopy
TEM: transmission electron microscopy
UC: ultracentrifugation
%CV: coefficient of variance

## I. Ethical approval

Not applicable

## J. Consent for publications

All co-authors have consented to publication

## K. Data availability

The mass spectrometry proteomics data have been deposited to the ProteomeXchange Consortium via the PRIDE partner repository with the dataset identifiers PXD051938 (comparison FAEVEr with UC), PXD051955 (comparison MCF7 proteome of EVs and cells under starving conditions) and PXD051956 (comparison FBS with EDS). All required details concerning the enrichment of extracellular vesicles were submitted to EV-TRACK (EV240045).

## N. Author contributions

JP planned, designed and performed the experiments and wrote the manuscript. JVDV, FDM and DDP assisted with the experiments. TVDS performed the HEK293T cell culture and production of rEV material. FB performed the electron microscopy. All authors read and approved the manuscript.

## Acknowledgements

The authors thank all operators of the VIB Proteomics Core for their excellence and input. The authors thank the lab of An Hendrix for kindly providing us with the required purified rEV aliquots.

## References

1. Dixson AC, Dawson TR, Di Vizio D, Weaver AM. Context-specific regulation of extracellular vesicle biogenesis and cargo selection. Nat Rev Mol Cell Biol. 2023 Jul 1;24(7):454–76.

2. van Niel G, D’Angelo G, Raposo G. Shedding light on the cell biology of extracellular vesicles. Nat Rev Mol Cell Biol. 2018 Apr 1;19(4):213–28.

3. Colombo M, Moita C, van Niel G, Kowal J, Vigneron J, Benaroch P, et al. Analysis of ESCRT functions in exosome biogenesis, composition and secretion highlights the heterogeneity of extracellular vesicles. J CELL Sci. 2013 Dec 15;126(24):5553–65.

4. Wei D, Zhan W, Gao Y, Huang L, Gong R, Wang W, et al. RAB31 marks and controls an ESCRT- independent exosome pathway. Cell Res. 2021 Feb 1;31(2):157–77.

5. Mathieu M, Martin-Jaular L, Lavieu G, Théry C. Specificities of secretion and uptake of exosomes and other extracellular vesicles for cell-to-cell communication. Nat Cell Biol. 2019 Jan 1;21(1):9–17.

6. Yu L, Zhu G, Zhang Z, Yu Y, Zeng L, Xu Z, et al. Apoptotic bodies: bioactive treasure left behind by the dying cells with robust diagnostic and therapeutic application potentials. J Nanobiotechnology. 2023 Jul 12;21(1):218.

7. Théry C, Witwer KW, Aikawa E, Alcaraz MJ, Anderson JD, Andriantsitohaina R, et al. Minimal information for studies of extracellular vesicles 2018 (MISEV2018): a position statement of the International Society for Extracellular Vesicles and update of the MISEV2014 guidelines. J Extracell Vesicles. 2018 Dec 1;7(1):1535750.

8. Cocucci E, Meldolesi J. Ectosomes and exosomes: shedding the confusion between extracellular vesicles. Trends Cell Biol. 2015 Jun 1;25(6):364–72.

9. Hadisurya M, Lee ZC, Luo Z, Zhang G, Ding Y, Zhang H, et al. Data-Independent Acquisition Phosphoproteomics of Urinary Extracellular Vesicles Enables Renal Cell Carcinoma Grade Differentiation. Mol Cell Proteomics. 2023 May 1;22(5):100536.

10. Hoshino A, Costa-Silva B, Shen TL, Rodrigues G, Hashimoto A, Tesic Mark M, et al. Tumour exosome integrins determine organotropic metastasis. Nature. 2015 Nov 1;527(7578):329–35.

11. Joshi BS, de Beer MA, Giepmans BNG, Zuhorn IS. Endocytosis of Extracellular Vesicles and Release of Their Cargo from Endosomes. ACS Nano. 2020 Apr 28;14(4):4444–55.

12. Stahl PD, Raposo G. Extracellular Vesicles: Exosomes and Microvesicles, Integrators of Homeostasis. Physiology. 2019 May 1;34(3):169–77.

13. Takahashi A, Okada R, Nagao K, Kawamata Y, Hanyu A, Yoshimoto S, et al. Exosomes maintain cellular homeostasis by excreting harmful DNA from cells. Nat Commun. 2017 May 16;8(1):15287.

14. Amin S, Massoumi H, Tewari D, Roy A, Chaudhuri M, Jazayerli C, et al. Cell Type-Specific Extracellular Vesicles and Their Impact on Health and Disease. Int J Mol Sci. 2024;25(5).

15. Hánělová K, Raudenská M, Masařík M, Balvan J. Protein cargo in extracellular vesicles as the key mediator in the progression of cancer. Cell Commun Signal. 2024 Jan 10;22(1):25.

16. Hoffmann O, Wormland S, Bittner AK, Collenburg M, Horn PA, Kimmig R, et al. Programmed death receptor ligand-2 (PD-L2) bearing extracellular vesicles as a new biomarker to identify early triple-negative breast cancer patients at high risk for relapse. J Cancer Res Clin Oncol. 2023 Mar 1;149(3):1159–74.

17. Zhang XW, Qi GX, Liu MX, Yang YF, Wang JH, Yu YL, et al. Deep Learning Promotes Profiling of Multiple miRNAs in Single Extracellular Vesicles for Cancer Diagnosis. ACS Sens [Internet]. 2024 Mar 5; Available from: 10.1021/acssensors.3c02789

18. Dorado E, Doria ML, Nagelkerke A, McKenzie JS, Maneta-Stavrakaki S, Whittaker TE, et al. Extracellular vesicles as a promising source of lipid biomarkers for breast cancer detection in blood plasma. J Extracell Vesicles. 2024 Mar 1;13(3):e12419.

19. Ma D, Xie A, Lv J, Min X, Zhang X, Zhou Q, et al. Engineered extracellular vesicles enable high- efficient delivery of intracellular therapeutic proteins. Protein Cell. 2024 Mar 22;pwae015.

20. Attem J, Narayana RVL, Manukonda R, Kaliki S, Vemuganti GK. Small extracellular vesicles loaded with carboplatin effectively enhance the cytotoxicity of drug-resistant cells from Y79 cells-in vitro. Biomed Pharmacother. 2024 Apr 1;173:116403.

21. Lone SN, Nisar S, Masoodi T, Singh M, Rizwan A, Hashem S, et al. Liquid biopsy: a step closer to transform diagnosis, prognosis and future of cancer treatments. Mol Cancer. 2022 Mar 18;21(1):79.

22. K S, T D, M P. Small extracellular vesicles as a multicomponent biomarker platform in urinary tract carcinomas. Front Mol Biosci [Internet]. 2022;9. Available from: https://www.frontiersin.org/articles/10.3389/fmolb.2022.916666

23. Gao X, Gao B, Li S. Extracellular vesicles: A new diagnostic biomarker and targeted drug in osteosarcoma. Front Immunol [Internet]. 2022;13. Available from: https://api.semanticscholar.org/CorpusID:252468251

24. Hirschberg Y, Valle-Tamayo N, Dols-Icardo O, Engelborghs S, Buelens B, Vandenbroucke RE, et al. Proteomic comparison between non-purified cerebrospinal fluid and cerebrospinal fluid-derived extracellular vesicles from patients with Alzheimer’s, Parkinson’s and Lewy body dementia. J Extracell Vesicles. 2023 Dec 1;12(12):12383.

25. Vandendriessche C, Kapogiannis D, Vandenbroucke RE. Biomarker and therapeutic potential of peripheral extracellular vesicles in Alzheimer’s disease. Adv Drug Deliv Rev. 2022 Nov 1;190:114486.

26. Newman LA, Muller K, Rowland A. Circulating cell-specific extracellular vesicles as biomarkers for the diagnosis and monitoring of chronic liver diseases. Cell Mol Life Sci. 2022 Apr 10;79(5):232.

27. Mazzucco M, Mannheim W, Shetty SV, Linden JR. CNS endothelial derived extracellular vesicles are biomarkers of active disease in multiple sclerosis. Fluids Barriers CNS. 2022 Feb 8;19(1):13.

28. Horvatovich PL, Bischoff R. Current Technological Challenges in Biomarker Discovery and Validation. Eur J Mass Spectrom. 2010 Feb 1;16(1):101–21.

29. Schiess R, Wollscheid B, Aebersold R. Targeted proteomic strategy for clinical biomarker discovery. Mol Oncol. 2009 Feb 1;3(1):33–44.

30. Burton JB, Carruthers NJ, Stemmer PM. Enriching extracellular vesicles for mass spectrometry. Mass Spectrom Rev. 2023 Mar 1;42(2):779–95.

31. Demichev V, Messner CB, Vernardis SI, Lilley KS, Ralser M. DIA-NN: neural networks and interference correction enable deep proteome coverage in high throughput. Nat Methods. 2020 Jan 1;17(1):41–4.

32. Gessulat S, Schmidt T, Zolg DP, Samaras P, Schnatbaum K, Zerweck J, et al. Prosit: proteome-wide prediction of peptide tandem mass spectra by deep learning. Nat Methods. 2019 Jun 1;16(6):509–18.

33. Guzman UH, Martinez-Val A, Ye Z, Damoc E, Arrey TN, Pashkova A, et al. Ultra-fast label- free quantification and comprehensive proteome coverage with narrow-window data- independent acquisition. Nat Biotechnol [Internet]. 2024 Feb 1; Available from: 10.1038/s41587-023-02099-7

34. Yoshitake J, Azami M, Sei H, Onoshima D, Takahashi K, Hirayama A, et al. Rapid Isolation of Extracellular Vesicles Using a Hydrophilic Porous Silica Gel-Based Size-Exclusion Chromatography Column. Anal Chem. 2022 Oct 11;94(40):13676–81.

35. Vanderboom PM, Dasari S, Ruegsegger GN, Pataky MW, Lucien F, Heppelmann CJ, et al. A size-exclusion-based approach for purifying extracellular vesicles from human plasma. Cell Rep Methods. 2021 Jul 26;1(3):100055.

36. Onódi Z, Pelyhe C, Terézia Nagy C, Brenner GB, Almási L, Kittel Á, et al. Isolation of High- Purity Extracellular Vesicles by the Combination of Iodixanol Density Gradient Ultracentrifugation and Bind-Elute Chromatography From Blood Plasma. Front Physiol [Internet]. 2018;9. Available from: https://www.frontiersin.org/journals/physiology/articles/10.3389/fphys.2018.01479

37. Gallart-Palau X, Serra A, Wong ASW, Sandin S, Lai MKP, Chen CP, et al. Extracellular vesicles are rapidly purified from human plasma by PRotein Organic Solvent PRecipitation (PROSPR). Sci Rep. 2015 Sep 30;5(1):14664.

38. Rider MA, Hurwitz SN, Meckes DG. ExtraPEG: A Polyethylene Glycol-Based Method for Enrichment of Extracellular Vesicles. Sci Rep. 2016 Apr 12;6(1):23978.

39. Brambilla D, Sola L, Ferretti AM, Chiodi E, Zarovni N, Fortunato D, et al. EV Separation: Release of Intact Extracellular Vesicles Immunocaptured on Magnetic Particles. Anal Chem. 2021 Apr 6;93(13):5476–83.

40. Vergauwen G, Tulkens J, Pinheiro C, Avila Cobos F, Dedeyne S, De Scheerder MA, et al. Robust sequential biophysical fractionation of blood plasma to study variations in the biomolecular landscape of systemically circulating extracellular vesicles across clinical conditions. J Extracell Vesicles. 2021 Aug 1;10(10):e12122.

41. Van Dorpe S, Lippens L, Boiy R, Pinheiro C, Vergauwen G, Rappu P, et al. Integrating automated liquid handling in the separation workflow of extracellular vesicles enhances specificity and reproducibility. J Nanobiotechnology. 2023 May 19;21(1):157.

42. An M, Wu J, Zhu J, Lubman DM. Comparison of an Optimized Ultracentrifugation Method versus Size-Exclusion Chromatography for Isolation of Exosomes from Human Serum. J Proteome Res. 2018 Oct 5;17(10):3599–605.

43. Turner NP, Abeysinghe P, Kwan Cheung KA, Vaswani K, Logan J, Sadowski P, et al. A Comparison of Blood Plasma Small Extracellular Vesicle Enrichment Strategies for Proteomic Analysis. Proteomes. 2022;10(2).

44. Brennan K, Martin K, FitzGerald SP, O’Sullivan J, Wu Y, Blanco A, et al. A comparison of methods for the isolation and separation of extracellular vesicles from protein and lipid particles in human serum. Sci Rep. 2020 Jan 23;10(1):1039.

45. Baranyai T, Herczeg K, Onódi Z, Voszka I, Módos K, Marton N, et al. Isolation of Exosomes from Blood Plasma: Qualitative and Quantitative Comparison of Ultracentrifugation and Size Exclusion Chromatography Methods. PLOS ONE. 2015 Dec 21;10(12):e0145686.

46. Askeland A, Borup A, Østergaard O, Olsen JV, Lund SM, Christiansen G, et al. Mass-Spectrometry Based Proteome Comparison of Extracellular Vesicle Isolation Methods: Comparison of ME-kit, Size-Exclusion Chromatography, and High-Speed Centrifugation. Biomedicines. 2020;8(8).

47. Van Deun J, Mestdagh P, Agostinis P, Akay Ö, Anand S, Anckaert J, et al. EV-TRACK: transparent reporting and centralizing knowledge in extracellular vesicle research. Nat Methods. 2017 Mar 1;14(3):228–32.

48. Geeurickx E, Lippens L, Rappu P, De Geest BG, De Wever O, Hendrix A. Recombinant extracellular vesicles as biological reference material for method development, data normalization and assessment of (pre-)analytical variables. Nat Protoc. 2021 Feb 1;16(2):603– 33.

49. Geeurickx E, Tulkens J, Dhondt B, Van Deun J, Lippens L, Vergauwen G, et al. The generation and use of recombinant extracellular vesicles as biological reference material. Nat Commun. 2019 Jul 23;10(1):3288.

50. Sidhom K, Obi PO, Saleem A. A Review of Exosomal Isolation Methods: Is Size Exclusion Chromatography the Best Option? Int J Mol Sci. 2020;21(18).

51. Ding M, Wang C, Lu X, Zhang C, Zhou Z, Chen X, et al. Comparison of commercial exosome isolation kits for circulating exosomal microRNA profiling. Anal Bioanal Chem. 2018 Jun 1;410(16):3805–14.

52. Royo F, Zuñiga-Garcia P, Sanchez-Mosquera P, Egia A, Perez A, Loizaga A, et al. Different EV enrichment methods suitable for clinical settings yield different subpopulations of urinary extracellular vesicles from human samples. J Extracell Vesicles. 2016 Jan 1;5(1):29497.

53. Lobb RJ, Becker M, Wen Wen S, Wong CSF, Wiegmans AP, Leimgruber A, et al. Optimized exosome isolation protocol for cell culture supernatant and human plasma. J Extracell Vesicles. 2015 Jan 1;4(1):27031.

54. Guan S, Yu H, Yan G, Gao M, Sun W, Zhang X. Characterization of Urinary Exosomes Purified with Size Exclusion Chromatography and Ultracentrifugation. J Proteome Res. 2020 Jun 5;19(6):2217–25.

55. Gámez-Valero A, Monguió-Tortajada M, Carreras-Planella L, Franquesa M, Beyer K, Borràs FE. Size-Exclusion Chromatography-based isolation minimally alters Extracellular Vesicles’ characteristics compared to precipitating agents. Sci Rep. 2016 Sep 19;6(1):33641.

56. Vergauwen G, Dhondt B, Van Deun J, De Smedt E, Berx G, Timmerman E, et al. Confounding factors of ultrafiltration and protein analysis in extracellular vesicle research. Sci Rep. 2017 Jun 2;7(1):2704.

57. Jordan KR, Hall JK, Schedin T, Borakove M, Xian JJ, Dzieciatkowska M, et al. Extracellular vesicles from young women’s breast cancer patients drive increased invasion of non-malignant cells via the Focal Adhesion Kinase pathway: a proteomic approach. Breast Cancer Res. 2020 Nov 23;22(1):128.

58. Allelein S, Medina-Perez P, Lopes ALH, Rau S, Hause G, Kölsch A, et al. Potential and challenges of specifically isolating extracellular vesicles from heterogeneous populations. Sci Rep. 2021 Jun 2;11(1):11585.

59. Eyckerman S, Titeca K, Van Quickelberghe E, Cloots E, Verhee A, Samyn N, et al. Trapping mammalian protein complexes in viral particles. Nat Commun. 2016 Apr 28;7(1):11416.

60. Wang J, Vasaikar S, Shi Z, Greer M, Zhang B. WebGestalt 2017: a more comprehensive, powerful, flexible and interactive gene set enrichment analysis toolkit. Nucleic Acids Res. 2017 Jul 3;45(W1):W130–7.

61. HaileMariam M, Eguez RV, Singh H, Bekele S, Ameni G, Pieper R, et al. S-Trap, an Ultrafast Sample-Preparation Approach for Shotgun Proteomics. J Proteome Res. 2018 Sep 7;17(9):2917–24.

62. Ono A. HIV-1 assembly at the plasma membrane: Gag trafficking and localization. Future Virol. 2009 May 1;4(3):241–57.

63. Chhuon C, Zhang SY, Jung V, Lewandowski D, Lipecka J, Pawlak A, et al. A sensitive S-Trap-based approach to the analysis of T cell lipid raft proteome. J Lipid Res. 2020 Nov 1;61(11):1512–23.

64. Elinger D, Gabashvili A, Levin Y. Suspension Trapping (S-Trap) Is Compatible with Typical Protein Extraction Buffers and Detergents for Bottom-Up Proteomics. J Proteome Res. 2019 Mar 1;18(3):1441–5.

65. Ludwig KR, Schroll MM, Hummon AB. Comparison of In-Solution, FASP, and S-Trap Based Digestion Methods for Bottom-Up Proteomic Studies. J Proteome Res. 2018 Jul 6;17(7):2480–90.

66. Varnavides G, Madern M, Anrather D, Hartl N, Reiter W, Hartl M. In Search of a Universal Method: A Comparative Survey of Bottom-Up Proteomics Sample Preparation Methods. J Proteome Res. 2022 Oct 7;21(10):2397–411.

67. Osteikoetxea X, Sódar B, Németh A, Szabó-Taylor K, Pálóczi K, Vukman KV, et al. Differential detergent sensitivity of extracellular vesicle subpopulations. Org Biomol Chem. 2015;13(38):9775–82.

68. Osteikoetxea X, Balogh A, Szabó-Taylor K, Németh A, Szabó TG, Pálóczi K, et al. Improved Characterization of EV Preparations Based on Protein to Lipid Ratio and Lipid Properties. PLOS ONE. 2015 Mar 23;10(3):e0121184.

69. Choi DS, Kim DK, Kim YK, Gho YS. Proteomics, transcriptomics and lipidomics of exosomes and ectosomes. PROTEOMICS. 2013 May 1;13(10–11):1554–71.

70. Subra C, Laulagnier K, Perret B, Record M. Exosome lipidomics unravels lipid sorting at the level of multivesicular bodies. Recent Dev Biosynth Storage Transp Lipids. 2007 Feb 1;89(2):205–12.

71. Minhua Feng, Berdugo Morales A, Poot A, Beugeling T, Bantjes A. Effects of Tween 20 on the desorption of proteins from polymer surfaces. J Biomater Sci Polym Ed. 1996 Jan 1;7(5):415– 24.

72. Amirilargani M, Saljoughi E, Mohammadi T. Improvement of permeation performance of polyethersulfone (PES) ultrafiltration membranes via addition of Tween-20. J Appl Polym Sci. 2010 Jan 5;115(1):504–13.

73. Lee KJ, Mower R, Hollenbeck T, Castelo J, Johnson N, Gordon P, et al. Modulation of Nonspecific Binding in Ultrafiltration Protein Binding Studies. Pharm Res. 2003 Jul 1;20(7):1015–21.

74. Urzì O, Olofsson Bagge R, Crescitelli R. The dark side of foetal bovine serum in extracellular vesicle studies. J Extracell Vesicles. 2022 Oct 1;11(10):e12271.

75. Drabovich AP, Pavlou MP, Dimitromanolakis A, Diamandis EP. Quantitative Analysis of Energy Metabolic Pathways in MCF-7 Breast Cancer Cells by Selected Reaction Monitoring Assay *. Mol Cell Proteomics. 2012 Aug 1;11(8):422–34.

76. Nie S, McDermott SP, Deol Y, Tan Z, Wicha MS, Lubman DM. A quantitative proteomics analysis of MCF7 breast cancer stem and progenitor cell populations. PROTEOMICS. 2015 Nov 1;15(22):3772–83.

77. Leidal AM, Huang HH, Marsh T, Solvik T, Zhang D, Ye J, et al. The LC3-conjugation machinery specifies the loading of RNA-binding proteins into extracellular vesicles. Nat Cell Biol. 2020 Feb 1;22(2):187–99.

78. Fujita N, Itoh T, Omori H, Fukuda M, Noda T, Yoshimori T. The Atg16L Complex Specifies the Site of LC3 Lipidation for Membrane Biogenesis in Autophagy. Mol Biol Cell. 2008 May 1;19(5):2092–100.

79. Matsushita M, Suzuki NN, Obara K, Fujioka Y, Ohsumi Y, Inagaki F. Structure of Atg5·Atg16, a Complex Essential for Autophagy *. J Biol Chem. 2007 Mar 2;282(9):6763–72.

80. Urbańska K, Orzechowski A. The Secrets of Alternative Autophagy. Cells. 2021;10(11).

81. Salimi L, Akbari A, Jabbari N, Mojarad B, Vahhabi A, Szafert S, et al. Synergies in exosomes and autophagy pathways for cellular homeostasis and metastasis of tumor cells. Cell Biosci. 2020 May 13;10(1):64.

82. Pirkmajer S, Chibalin AV. Serum starvation: caveat emptor. Am J Physiol-Cell Physiol. 2011 Aug 1;301(2):C272–9.

83. Shin J, Rhim J, Kwon Y, Choi SY, Shin S, Ha CW, et al. Comparative analysis of differentially secreted proteins in serum-free and serum-containing media by using BONCAT and pulsed SILAC. Sci Rep. 2019 Feb 28;9(1):3096.

