## Supplementary material for "Filter-aided extracellular vesicle enrichment (FAEVEr) for proteomics": Suppl figures

Supplementary figure 01

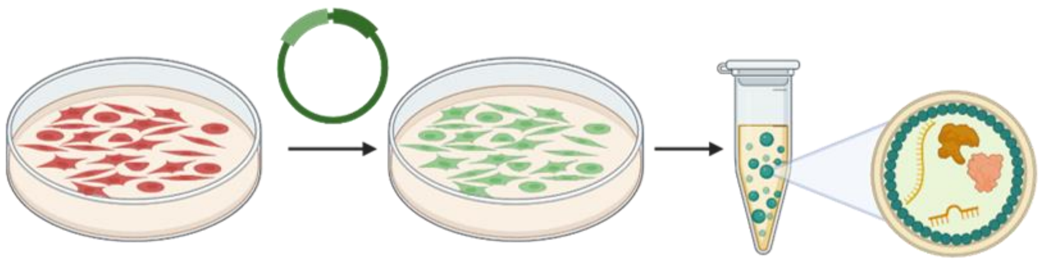

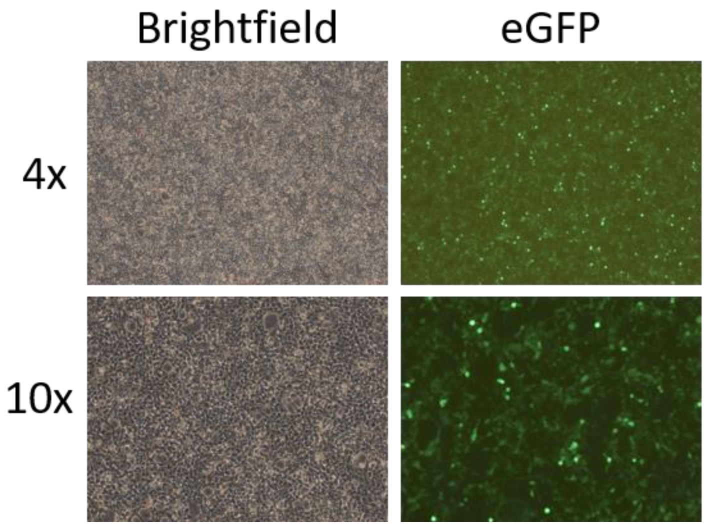

Supplementary figure 02

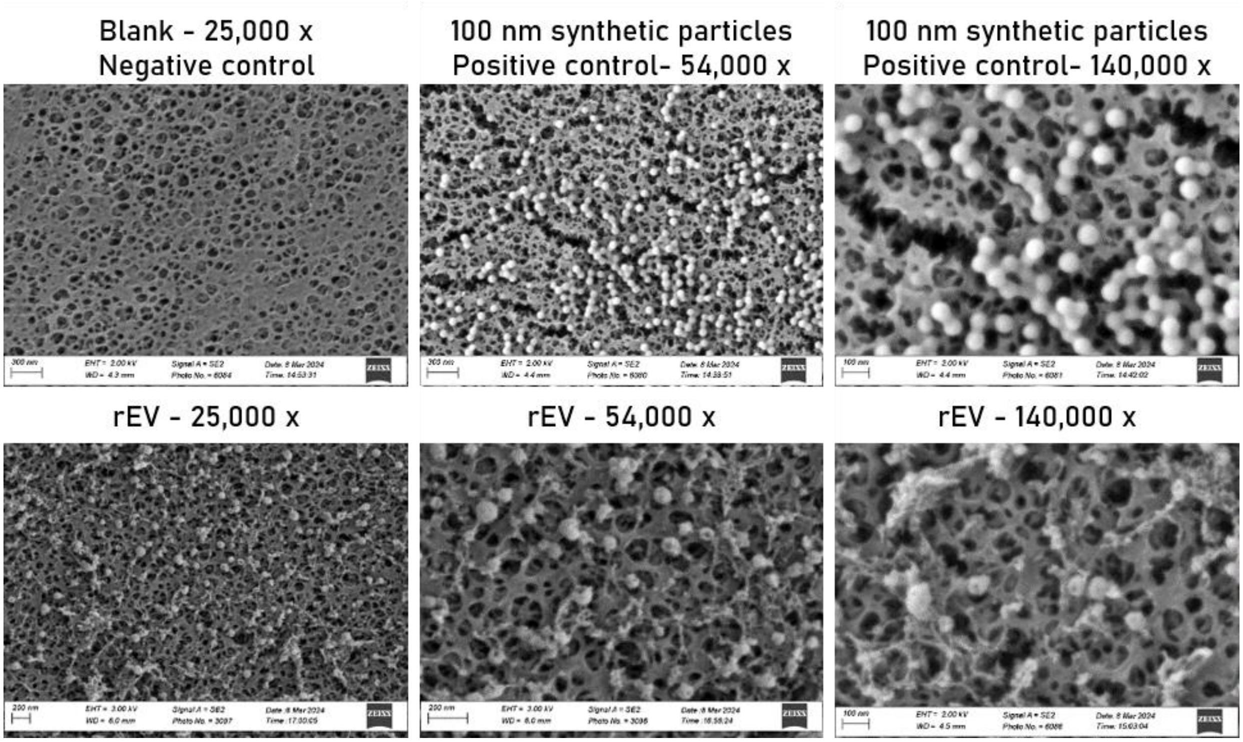

Supplementary figure 03

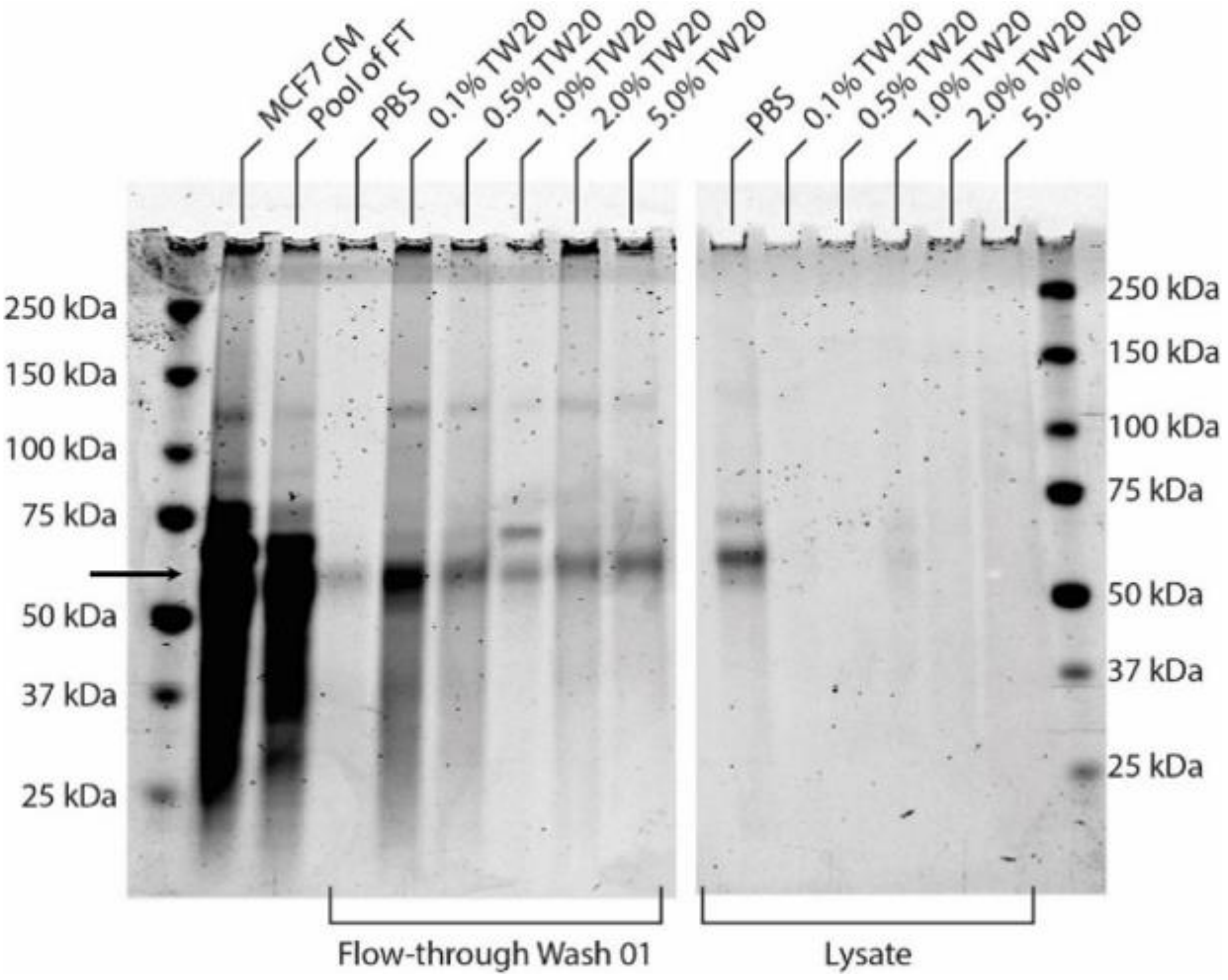

Supplementary figure 04

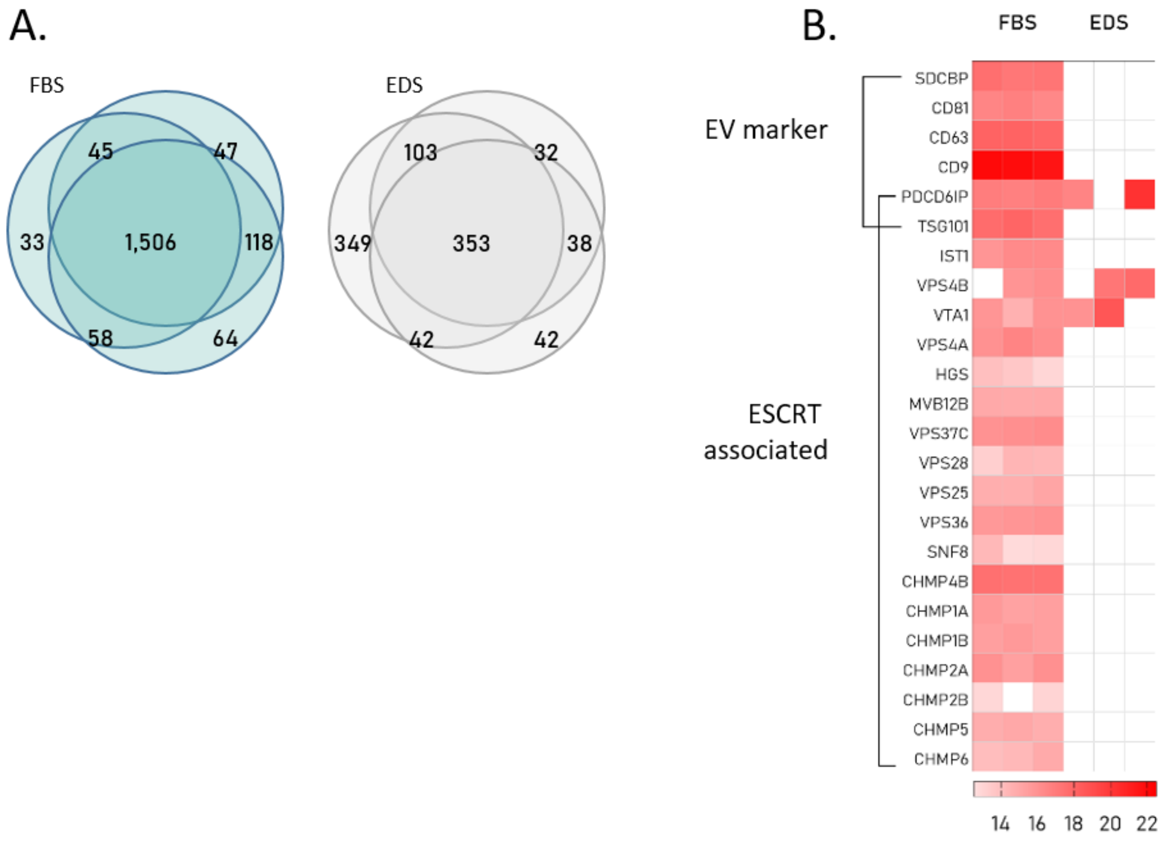

Supplementary figure 05

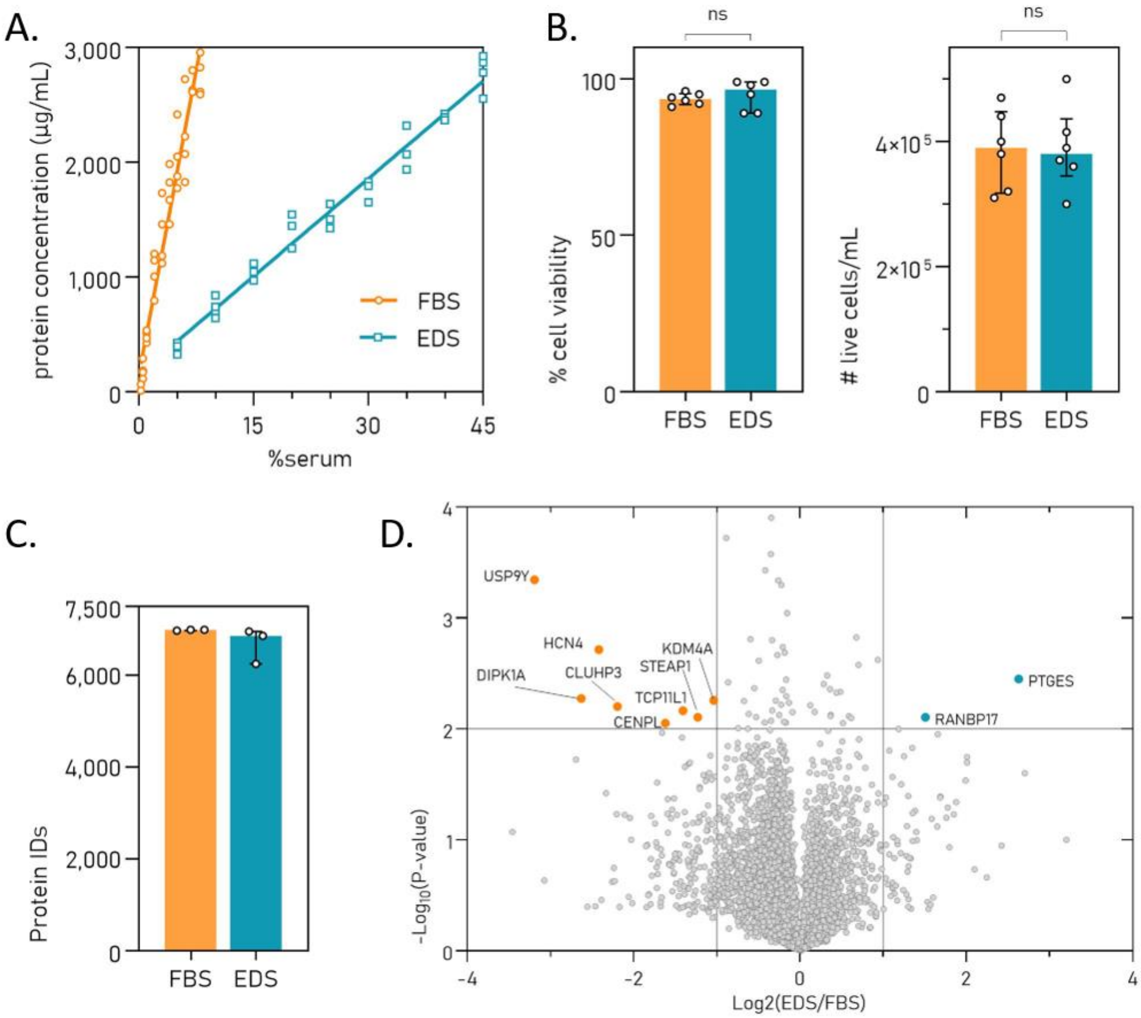

Supplementary figure 06

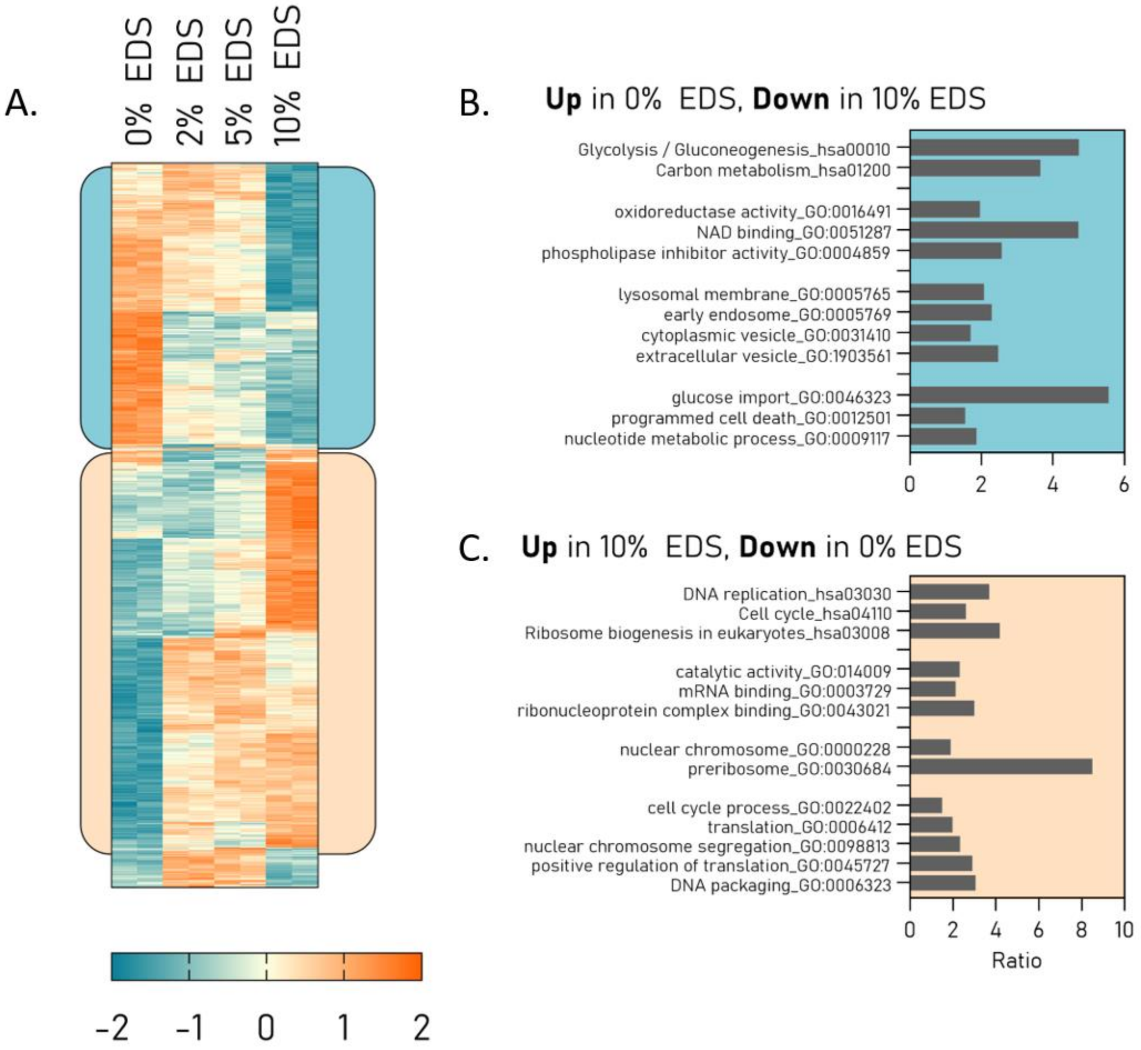
